# Incongruent end structures of leading and lagging telomeres dictate the nature of end replication problem

**DOI:** 10.1101/2025.02.23.639772

**Authors:** Tiantian Ye, Qingqing Yuan, Shuheng Wu, Jing-Tong Zhao, Zhi-Jing Wu, Jia-Cheng Liu, Wei Wu, Jin-Qiu Zhou

## Abstract

The end replication problem refers to the incomplete replication of parental DNA at telomeres, a process whose molecular depiction is hampered by the complex nature of telomere ends. Here we recapitulate this process using a synthetic *de novo* telomere in yeast and delineate distinct molecular fates of telomere ends *in vivo*. We show that the lagging strand telomere carries a ∼10 nt 3’ overhang, while the leading strand telomere has a Yku-protected blunt end, which is prevalent on native telomeres. In addition, RNase H2 but not RNase H1 is mainly responsible for the removal of the last RNA primer. Consistently, in the absence of RNase H2 activity, RNA primer is retained on the lagging strand telomere, attenuating telomere erosion and delaying senescence in telomerase-null cells. These findings highlight incongruent end structures on telomeres and clarify that the primary culprit behind end replication problem is the incompletely replicated lagging strand telomere.

## INTRODUCTION

Telomeres are the nucleoprotein structures at the ends of eukaryotic chromosomes, and protect chromosome ends from nucleolytic degradation and chromosome fusion^1,2^. In the budding yeast *Saccharomyces cerevisiae*, telomeric DNA spans approximately 300 base pairs and features a consensus sequence of TG_1-3_/C_1-3_A. Its protruding G-rich 3’ overhang typically measures between 12 to 15 nucleotides in length^3,4^. Many proteins bind at telomeres, for example, Rap1 directly binds to telomeric dsDNA, and recruits Rif1 and Rif2 to telomeres to negatively regulate telomere length^5,6^. Cdc13 binds to the the G-tail, and recruits Stn1 and Ten1 to protect telomere DNA from degradation^7^. Yku70/80 heterodimer binds the junction of dsDNA and ssDNA to protect telomeres^8^. These proteins function together to protect telomeres from being recognized as double-strand breaks and recruit telomerase and/or polymerase to elongate telomeres^2,9^.

Semiconservative DNA replication of a telomere generates two daughter telomeres (Figure S1)^4,10^. One is produced through leading-strand synthesis, and the other is produced through lagging-strand synthesis, also referred to as the leading strand and the lagging strand telomeres, respectively. The lagging strand telomere processing involves the removal of the ultimate RNA primer, which causes incomplete DNA replication of the parental DNA template at each round of DNA replication (Figure S1). Thus, telomeres shorten at each round of DNA replication, and eventually become critically short to trigger cellular senescence, known as the’end replication problem’^10–12^.

To overcome the end replication problem, most eukaryotes have evolved two pathways to lengthen telomeres, the telomerase pathway and the homologous recombination (HR) pathway^12,13^. Telomerase is a specialized reverse transcriptase that uses its intrinsic RNA as a template to elongate telomeric G-strand^11^. The complementary C-strand is synthesized by conventional DNA polymerases (i.e. Polα and δ)^14^. Thus, telomerase elongated telomeres eventually maintain a structure similar to the lagging strand telomere. In *S. cerevisiae*, telomerase consists of a number of subunits, including the catalytic subunit Est2, the RNA template Tlc1, the accessary subunits Est1 and Est3^2^. Est1 and Tlc1 directly interact with telomere-bound Cdc13 and Yku respectively to recruit telomerase^15^. In the absence of telomerase, homologous-recombination can be activated to back-up telomere lengthening^13^. Telomerase pathway appears to dominate over HR pathway in telomere lengthening as reactivation of telomerase effectively inhibits HR activity at telomeres^2,9^. Notably, both telomerase and HR prefer to elongate short telomeres, likely because long telomeres have an intact structure and are often overlooked by Mec1 or Tel1 kinases^2,16,17^.

As previous studies have constantly detected the 3’ overhangs on telomeres^3^, a fill-in model for leading-strand telomere replication has been proposed and widely accepted^4^. In this model, the blunt end product of leading strand is an intermediate, and has to be inevitably processed after replication. The processing includes 5’-end resection, fill-in synthesis and RNA primer removal, resulting in the formation of a protruding 3’ overhang, structurally identical to the lagging strand telomere (Figure S1)^2,4,10^. Accordingly, the replication of the C_1-3_A strand (CA strand) results in a leading telomere that is approximately 10-12 nucleotides shorter than the parental telomere. As a result, both leading strand and lagging strand contribute equally to the end replication problem. However, previous studies have consistently reported that the average telomere shortening rate in telomerase-null cells ranges from 2 to 3 base pairs per cell division^11,18^. This significant discrepancy between the proposed telomere shortening rate (∼5 bp, one-half of the length of 3’ overhang) and the actual experimental data (2-3 bp, one-quarter of the length of 3’ overhang) has led us to reconsider the primary cause of the end replication problem. It therefore seems imperative to dissect the exquisite molecular events involved in telomere replication.

During DNA replication, each Okazaki fragment synthesis initiate with a 7-12 nt RNA primer^19^. In the internal regions of a chromosome, the RNA primer is typically displaced by the adjacent Okazaki fragment to a 5’ flap and subsequently removed by endonucleases Rad27 (FEN1 in mammals) and/or Dna2^20^. At a telomere, however, this process differs because there is no upstream Okazaki fragment to extrude and displace the RNA primer at the very terminal Okazaki fragment in the lagging strand. Consequently, the terminal RNA primer cannot be removed by the same mechanism employed in the internal regions. So far, how the RNA primer at a telomere’s Okazaki fragment is removed remains mysterious. RNase H enzymes are endonucleases responsible for cleavage of ribo-nucleotide within an RNA-DNA hybrid. There are two types of RNase H enzymes, RNase H1 and RNase H2, both of which play critical roles in DNA-RNA hybrid removal and telomere regulation^21–23^. In the absence of both RNase Hs, the accumulation of telomeric RNA-DNA hybrid leads to delay of cellular senescence^24^, presumably by either preventing telomere shortening or promoting homologous recombination-mediated telomere extension, or both^24,25^. Although RNase H enzymes were reported to hydrolysis RNA primer *in vitro*^26,27^, whether it functions at last RNA primer removal *in vivo* remains elusive.

In this work, taking advantage of the *de novo* telomere system, we examined the end structure of leading and lagging strand telomeres in dividing yeast cells. We found that the lagging strand telomere carries a ∼10 nt 3’ overhang, but the leading strand telomere has a Yku-protected stable blunt end, contradicting to the widely accepted model that both leading and lagging strand telomeres contain an overhang^4^. In addition, with the high-resolution assay developed in this work, we have identified for the first time that nuclease RNase H2 is responsible for the removal of the last RNA primer. Furthermore, RNA primer retention at the lagging strand telomere attenuates telomere erosion and delays senescence in a TERRA-independent manner. These results reveal incongruent end structures on leading and lagging strand telomeres, highlighting the unequal contributions of incompletely replicated lagging strand telomeres and fully replicated leading strand telomeres to the end replication problem.

## RESULTS

### *De novo* telomere with long telomeric sequence is refractory to end processing

In *S. cerevisiae*, a haploid cell possesses 32 telomeres, each varying in length both among cells and within the same cell. Telomere end processing after replication involves subtle nucleotide changes of up to 10-30 at chromosome ends, making it hard to monitor a single native telomere^28,29^. In addition, short telomeres are preferentially elongated by telomerase followed by fill-in synthesis^30^, further complicating the detection of telomere end processing. To dissect end processing events precisely, we utilized a long *de novo* telomere induction system, known for its sensitive readouts^31–33^. The engineered chromosome arm comprises a 250 bp TG_1-3_/C_1-3_A telomeric sequence flanked by a specific CRISPR-Cas9 gRNA targeting site and the *KanMX* gene. Adjacent to the gRNA recognition site, we designed a (TGG(TG)_5_)_2_ sequence to render it unreadable for Rap1, but readable for Cdc13 if the *de novo* telomere terminates with a 3’ overhang (Figure 1A)^34,35^. After galactose-induced gRNA expression, constitutively expressed Cas9 creates a double-strand break at the target site, resulting in the formation of *de novo* telomere with a blunt end (Figure 1A). Efficient induction of long *de novo* telomeres at G2/M phase were assessed via native Southern blotting (Figures S2A-S2D). The signal of terminal region of *de novo* telomeres (indicated by E-TRF: EcoR V digested terminal restriction fragment) became more intensive as the induction time increased (Figure S2D), consistent with the high stability of long *de novo* telomere, which is not a favorable target of telomerase ^32^.

**Figure 1.**
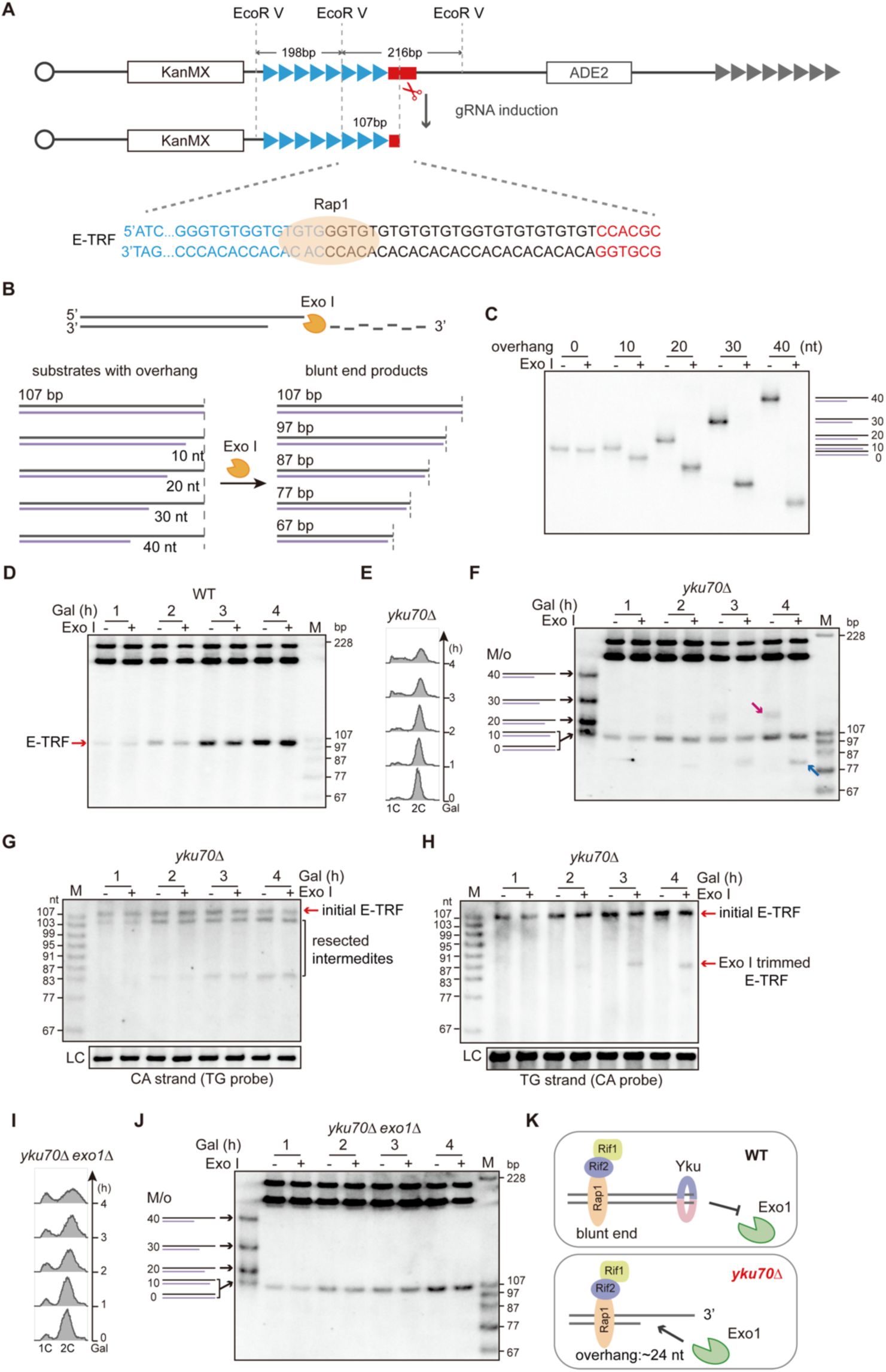
***De novo* telomere is capped by Yku, preventing from Exo1-mediated resection.** (A) Schematic illustration of the *de novo* telomere induction system (not to scale, EcoR V sites are displayed), a *KanMX*-250-bp TG_1-3_/C_1-3_A cassette is inserted in *ADH4* locus and an *ADE2* marker is inserted in *MNT2* locus to monitor leaky expression, red rectangle indicates gRNA recognition site, blue and grey triangles represent the inserted and native telomeric tracts, respectively. CRISPR-Cas9 cut results in the formation of *de novo* telomere. E-TRF sequence is shown at the bottom: sequence in blue, suitable for Rap1 binding, first Rap1 binding site is labeled; sequence in black, suitable for Cdc13 binding; sequence in red: non-TG sequence of Cas9 cleavage. (B) Schematic illustration of the synthesized oligos with 0-to 40-nt 3’ overhang, gray line: TG strand, purple line: CA strand. Exo I trims 3’ overhang from the direction of 3’ to 5’. Left: structures of oligos before Exo I digestion, the length of 3’ overhang is labeled under the TG strand. Right: structures of oligos after Exo I digestion, the length of double strand region is labeled on the top. (C) Southern blotting validation of Exo I treatment of synthesized oligos with 0-to 40-nt 3’ overhang. (D, F and J) Southern blotting detection of the 3’ overhang of the *de novo* telomere (E-TRF) induced in WT (D), *yku7*0Δ (F) and *yku70*Δ *exo1*Δ (J) cells at G2/M phase. Overhanged marker (M/o) are labeled on left. (E and I) FACS analysis of DNA content of nocodazole-synchronized *yku70*Δ (E) and *yku70*Δ *exo1*Δ (I) cells. (G and H) Denatured Southern blotting detection of the CA-strand (G) and TG-strand (H) in *yku7*0Δ cells, DNA markers (M) are labeled on left, LC: loading control. (K) Model for *de novo* telomere end protection and processing in WT and *yku70*Δ cells (see text for details). See also Figures S2 and S3.

To examine the end structure of *de novo* telomere in detail, we employed Exo I, a single-stranded DNA exonuclease of *E. coli,* which is widely used to remove the 3’ overhang at telomeric DNA ends *in vitro*^4,31^. Exo I activity was verified on various oligonucleotides by native or denatured Southern blotting (Figure 1B). When oligonucleotide samples were separated on native PAGE, both the length of the double-stranded DNA and the length of the 3’ overhang affected their mobility, i.e., the longer the 3’ overhangs, the slower the migration rate before Exo I digestion (Figure 1C). Exo I digestion of the overhangs accelerated their mobility to varying degrees, and each trimmed product migrated to a position well aligned with blunt-ended double-stranded oligos (Figure 1C), suggesting that Exo I digests the overhangs efficiently. Denatured Southern blotting confirmed that the CA strand is not affected by Exo I (Figure S2E), but the single-stranded TG-rich overhangs were efficiently trimmed by Exo I (Figure S2F), although measured a few nucleotides longer than their complementary CA-rich oligos (Figures S2E and S2F), which is because that short 3’ overhangs are not favorable substrates for Exo I. These oligonucleotides were used as markers in the following experiments.

When Exo I digestion was applied to the E-TRFs, the migration of the 107 bp band was not affected at any induction time point (Figure 1D), suggesting that the *de novo* telomere is blunt end and remains stable for at least 4 hours in G2/M phase. We also found that the *de novo* telomere formed in G1 phase behaved identically to that in G2/M phase (Figures S2G and S2H). These results indicate that *de novo* telomere is stable with a blunt end.

### Yku protects *de novo* telomere from Exo1-mediated resection

Telomeric TG_1-3_/C_1-3_A tracts are typically bound and safeguarded by Rap1, Rif1, Rif2 and Yku^5,8,36,37^. To investigate whether the maintenance of stable *de novo* telomere end requires Yku or Rif2, we assessed the end structure of *de novo* telomeres in the *yku70*Δ and *rif2*Δ cells arrested at G2/M phase. In *yku70*Δ cells (Figure 1E), a new band appeared, migrating more slowly than both the 107-bp band and the 20 nt overhanged marker (Figure 1F, magenta arrow), and its intensity increased during the 3-to 4-hour induction period; this band disappeared after *in vitro* Exo I treatment, and a distinct band of <87 bp appeared (Figure 1F, blue arrow). The slower migration and high sensitivity to Exo I digestion suggest that the *de novo* telomere in some of the *yku70*Δ cells carry a long 3’ overhang. Further denatured Southern blotting confirmed that a portion of the initial 107 nt CA strand were shortened to 103 and 83 nt (Figure 1G). TG strand was 107 nt before Exo I treatment, but part of the signal converted to 87 nt after Exo I treatment (Figure 1H). These results indicate that 5’ resection takes place at the CA-strand of the *de novo* telomere in *yku70*Δ cells, and the 3’ overhang, calculated from TG_(107 nt)_-CA_(83 nt)_, is ∼24 nt (Figures 1G and 1H). Given that most of the 107-bp signal in the native gel (Figure 1F) was resistant to Exo I digestion, we speculated that the 103-nt CA strand was likely generated through nicking by a cellular endonuclease (Figure 1G). Although previous studies showed that Rif2 and Rap1 play a major role in short *de novo* telomere protection^38,39^, we found that the *de novo* telomere induced in G2/M phase in *rif2*Δ cells was blunt ended (Figures S3A and S3B); and the *de novo* telomere end in the *yku70*Δ *rif2*Δ cells exhibited similar dynamics to those in *yku70*Δ cells (Figures S3C and S3D), indicating that Rif2 doesn’t play a major role in long telomere protection^40^. The preferential accumulation of the 83-nt band indicated a strictly controlled resection on the CA strand, which might be attributed to the involvement of other binding proteins. Since the Rap1 target sequence 5’-CACCCAC-3’ is positioned 28 nucleotides away from the end of the CA strand (Figure 1A), we hypothesized that Rap1 binding to the *de novo* telomere may act as a physical obstacle, preventing the CA-strand from extensive resection in *yku70*Δ cells.

Exo1, a 5’-3’ exonuclease, is responsible for ssDNA generation at telomeres in *yku70*Δ cells^41^. To investigate whether Exo1 involved in *de novo* telomere resection, we assessed the end structure of the *de novo* telomere in *exo1*Δ and *yku70*Δ *exo1*Δ mutants. In *exo1*Δ cells, the *de novo* telomere induced in G2/M phase exhibited blunt end structure, and the E-TRF was resistant to Exo I digestion (Figures S3E and S3F). In *yku70*Δ *exo1*Δ cells, the E-TRF of the uncapped *de novo* telomere remained stable during 4-hour induction and resistant to Exo I digestion (Figures 1I and 1J). Additional denatured Southern blotting showed that both the CA strand and the TG strand (Figures S3G and S3H) remained at their initial length of 107 nt, confirming no resection on CA strand. However, in *yku70*Δ cells, the absence of the Yku complex renders the telomere end susceptible to resection by Exo1, resulting in the generation of a 24 nt 3’ overhang (Figure 1K).

### The leading strand telomere maintains a stable blunt end after replication

We then monitored the end processing of *de novo* telomere-derived daughter telomeres after once replication, which presumably behave as native telomeres. Cells were first cultured with galactose to induce *de novo* telomere formation. Subsequently, they were arrested in early S phase with hydroxyurea (HU). Glucose was added to completely terminate gRNA expression. Finally, HU was washed out, and cells were released into the cell cycle with fresh glucose-containing media. Nocodazole was added after 60 minutes to prevent a second round of the cell cycle (Figure 2A). FACS analysis confirmed proper cell cycle progression as expected (Figure 2B). Prior to replication (Figure 2C, 0 min), the telomere end is blunt. After replication (Figure 2C, 120 min), the E-TRF migrated slightly slower (grey arrow), after Exo I digestion, two bands of 107 bp and 97 bp appeared (Figure 2C, indicated by blue and pink arrows, respectively). Importantly, the 107 bp band, which accounts for approximately half of the E-TRF signals, was still resistant to Exo I treatment (blue arrow).

**Figure 2.**
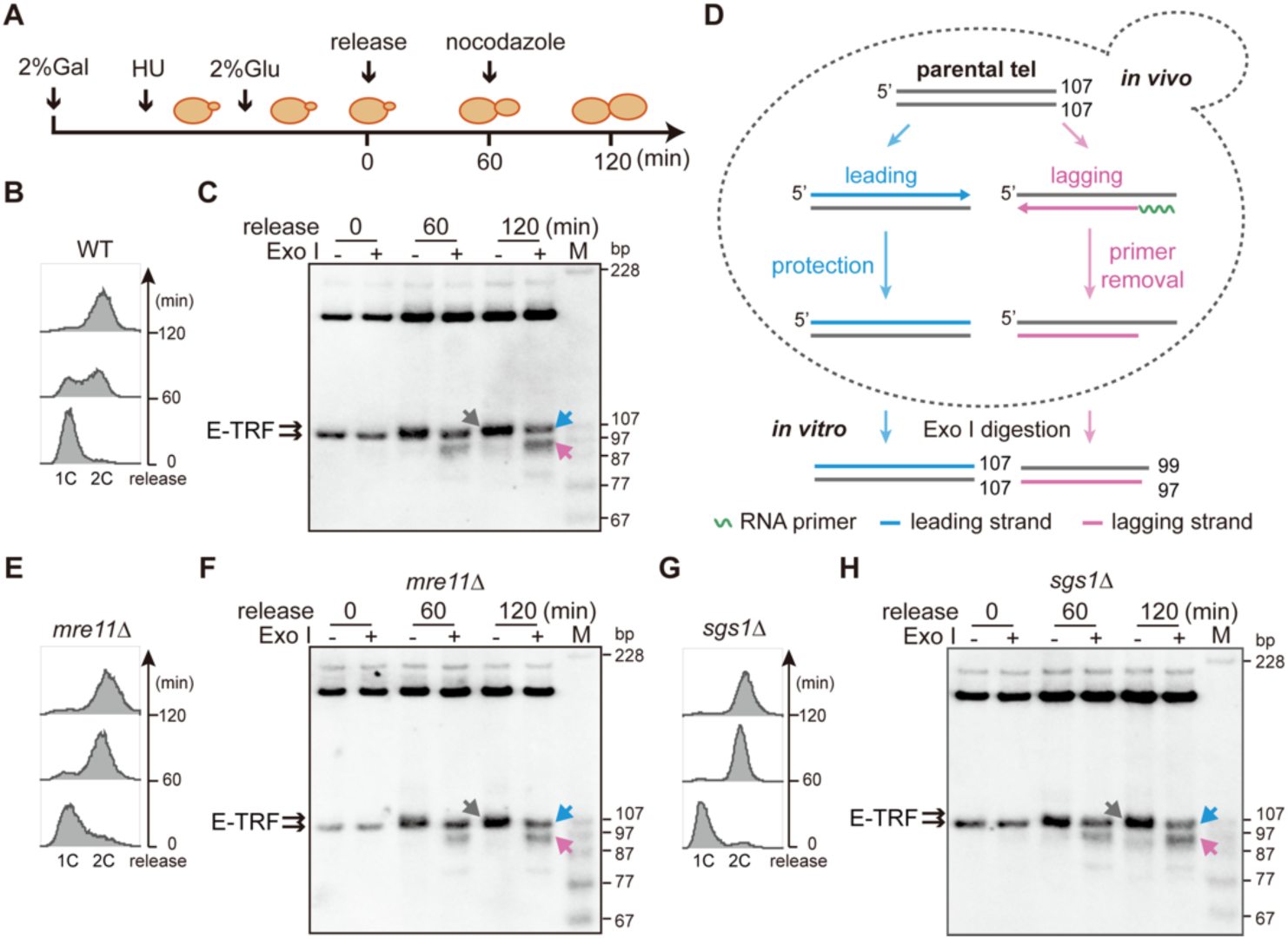
Half of the daughter telomeres are blunt-ended after DNA replication. (A) Flow chart of *de novo* telomere replication within one cell cycle (see text for detailed description). Cells are collected at the indicated time points for FACS and Southern blotting analysis. (B, E and G) FACS analysis of DNA content in WT (B), *mre11*Δ (E) and *sgs1*Δ (G) cells before and after release into cell cycle. (C, F and H) Southern blotting analysis of the 3’ overhang at both telomere ends during once DNA replication in WT (C), *mre11*Δ (F) and *sgs1*Δ (H) cells. Grey arrow: overlapped daughter telomeres, blue arrow: leading telomere, pink arrow: lagging telomere. (D) Schematic illustration of the E-TRF structures after DNA replication. See also Figure S4.

These results are contrary to the model that both daughter telomeres have a 3’ overhang (Figure S1)^4,10^, but fit the model that one of the daughter telomeres is blunt ended, the other of the daughter telomeres carries an ∼10 nt 3’ overhang (Figure S4A).

Further denatured Southern blotting showed precise lengths of each strand: before genome duplication, the CA strand was 107 nt (Figure S4B, 0 min); after replication, in addition to the initial 107 nt band, distinct bands of 99, 97, 95 and 83 nt emerged, which were supposed to be products left by *in vivo* RNA primer removal (Figure S4B, 120 min). The TG strand was 107 nt before genome duplication, and this signal increased after replication; whereas Exo I digestion decreased the signal approximately by half, converting to the bands of 101, 99, 97 and 87 nt (Figure S4C, 120 min). These results indicate that the 3’ overhang on the daughter telomere 2, calculated from TG_(107 nt)_-CA_(99/97/95 nt)_, is 8/10/12 nt (Figures S4A-S4C). Since the daughter telomeres we detected here included both the leading and lagging strand products, and the lagging strand must undergo RNA primer removal, our interpretations of these results are as follows.

(1) The signals that are sensitive to Exo I digestion originate from the lagging strand telomere, which possesses a 3’ overhang of ∼10 nt left by RNA primer removal. (2) The RNA priming by Polα-primase is initiated at/or near the 3’ end of the TG_1-3_ template strand, and the length of RNA primer is 8 to 12 nt long. (3) The signals that are resistant to Exo I treatment originate from the leading strand telomere, which has a blunt end (Figure 2D). Of note, a mild band of approximately 83 bp appeared after Exo I digestion (Figure 2C), suggesting that a small portion of telomere ends were inevitably not promptly and adequately protected by capping proteins (e.g., Yku).

To exclude the possibility that overhanged structure is resulted from the resection of leading strand telomere, we also detected the end structure in *mre11*Δ and *sae2*Δ cells, as both Mre11 and Sae2 have been reported to function in leading strand telomere resection^3,33,42^. Intriguingly, neither mutant exhibited defects in the generation of telomere 3’ overhang (∼10 nt) compared to WT cells (Figures 2E and 2F, S4D and S4E), even following RNase H treatment (Figure S4F), ruling out the potential effect of DNA-RNA hybrid on telomere end. Furthermore, inactivation of Sgs1, which is involved in long-range resection^43^, did not result in any changes in telomere end structure compared to WT cells either (Figures 2G and 2H). Thus, these findings consistently support the model that the leading strand telomere doesn’t undergo any processing, but rather remains blunt (Figure 2D).

### Blunt ends are prevalent on native telomeres

Although we unexpectedly observed the blunt-ended leading strand telomere, these results are based solely on the synthetic *de novo* telomere system. In fact, native telomeres were reported to constantly end with a 3’ overhang^3,9^. Therefore, it remains to be confirmed whether this blunt end also exists on native telomeres. Because the typical methods used for telomere examination are based on the structure of 3’ overhang^3,44,45^, which is not suitable for detecting blunt-ended telomeres. So, we utilized END-seq, an unbiased assay widely used to detect the ends of *in vivo* DNA double-strand breaks genome-wide^46–48^. To specifically investigate whether the blunt end is also a ubiquitous structure of native telomeres, we modified the typical END-seq protocol. As illustrated in the schematics, without any blunting treatment, the chromosomes embedded in agarose are directly subjected to the step of hairpin adaptor ligation (Figure 3A), which is essential for fragment enrichment and library preparation followed by next-generation sequencing (NGS). Thus, only native blunt ends can be captured in this way, sticky ends are excluded during the enrichment, hence referred to as native END-seq. Consequently, the sequencing reads starting with 5’ telomeric repeats (C_1-3_A) corresponding to blunt-ended native telomeres (Figure 3A). Consistent with the design, DNA ends with a 3’ overhang of 4 nt (generated by *in vitro* FseI digestion, Figure S5A) were barely detected in native END-seq but were significantly enriched in typical END-seq followed by end blunting (Figures S5B and S5C), indicating the blunt-end specificity of native END-seq. We then performed native END-seq in G2/M phase cells (Figure 3B), in which telomere replication is completed. Compared to other genomic loci, telomeric regions were highly enriched in WT cells (Figure 3C). These telomeric reads aligned well with the annotated telomeres via convergent telomeric sequence (Figures 3D and S6A), and disappeared when Exo III, a 3’ to 5’ exonuclease for blunt-ended double-stranded DNA, was used to digest the genome (Figure S6B). These results confirm that in addition to 3’ overhangs, blunt ends are another common feature of native telomeres.

**Figure 3.**
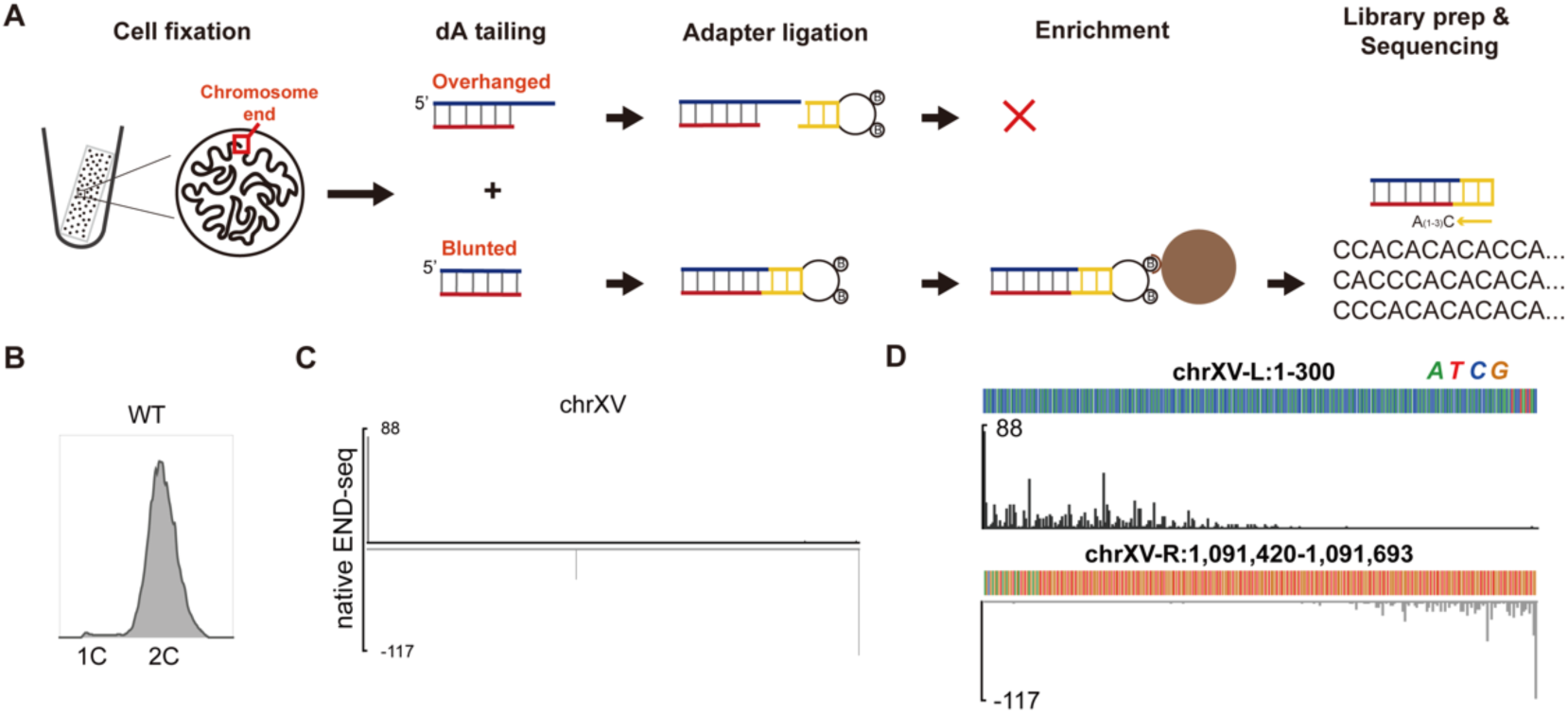
Blunt ends are readily detected on native telomeres. (A) Schematic illustration of the native END-seq procedures. (B) FACS analysis of DNA content in WT cells arrested by nocodazole. (C) Alignment of native END-seq reads detecting blunt DNA ends across chromosome XV in WT cells. Read coverage is colored in black or grey for plus and minus strand alignments, respectively. (D) Alignment of telomeric reads to annotated telomere of XV-L (top) and XV-R (bottom) in WT cells. Consistent with that native END-seq mainly captures C_(1-3)_A telomeric reads, telomeric reads from the left arm were aligned to the plus strand (black) and telomeric reads from right arm were aligned to the minus strand (grey). See also Figures S5 and S6.

### The blunt-ended leading-strand telomere is shielded by Yku

Since the Yku70/80 heterodimer caps the *de novo* telomere against Exo1 nuclease in G2/M phase (Figure 1K), and the leading strand telomere is structurally similar to G2/M phase induced *de novo* telomere, we thus hypothesized that Yku protect at leading telomere. To investigate the protective role of Yku at the blunt ended leading strand telomere, we analyzed telomere end structures in *yku70*Δ cells (Figures 4A and 4B). Before replication, a portion of telomere was resected, so, the E-TRF of the parental telomeres exhibited two main types: one measuring 107 bp with blunt end, the other carrying a 24 nt 3’ overhang (Figures 4B and S7A, 0 min). When replication was completed (Figure 4B, 120 min), the 107 bp band gradually diminished, while the smeared signals became increasingly prominent (Figure 4B, magenta arrow, 120 min). These signals represented the de-capped daughter telomeres including leading 1, lagging 1 and lagging 2 (Figure 4C), which contain a >20 nt 3’ overhang after *in vivo* 5’ resection, as they were sensitive to Exo I treatment, and converted to a <87 bp band (Figure 4B, yellow arrow). A small fraction of daughter telomeres showed an 83-bp E-TRF (Figure 4B, blue arrow), which was the product of daughter leading 2 (Figure 4C). It was resistant to Exo I treatment, suggesting that daughter leading 2 telomere had a blunt end and protected by Rap1 (Figure 1A).

**Figure 4.**
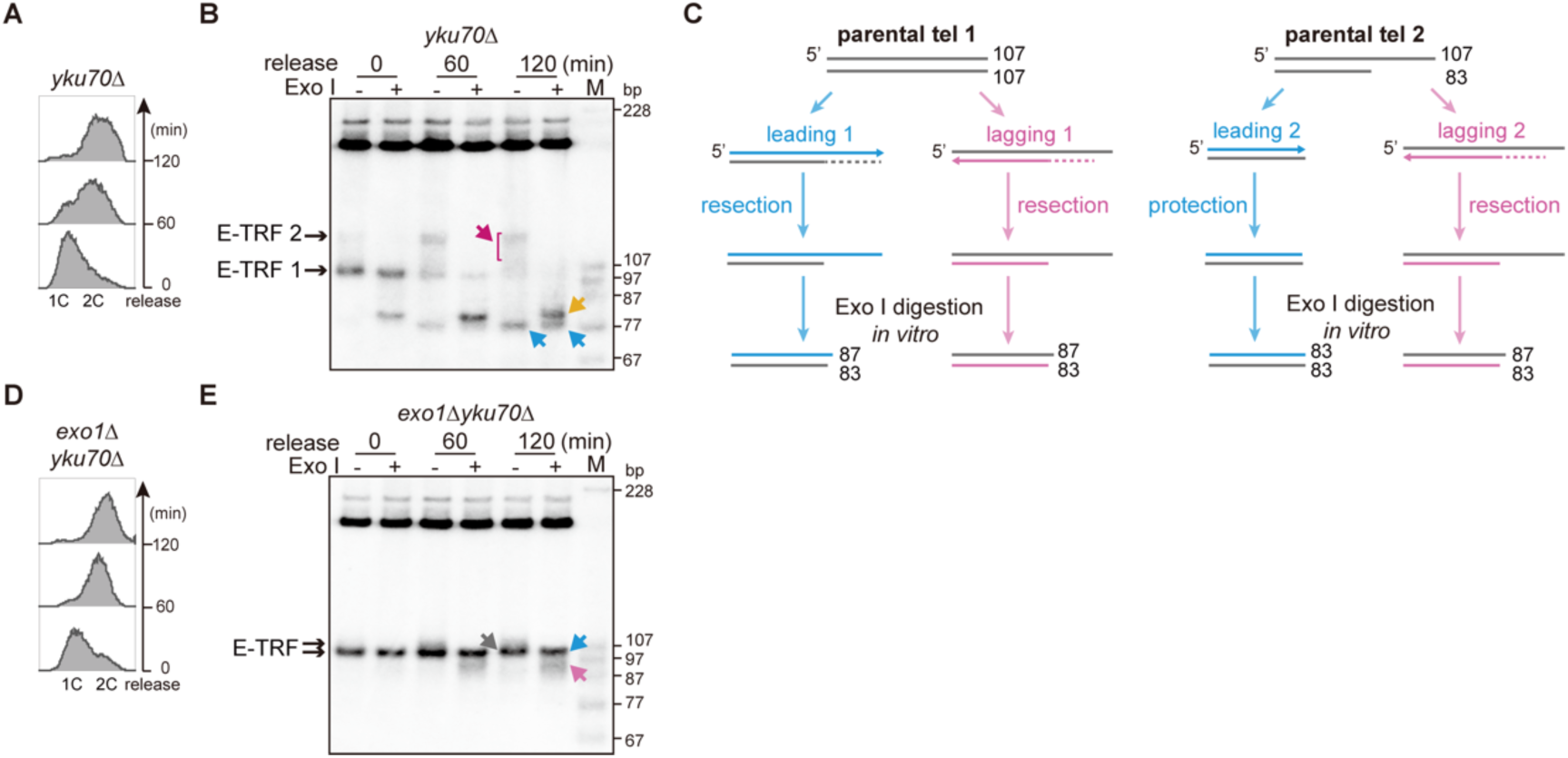
Both daughter telomeres are protected by Yku70. (A and D) FACS analysis of DNA content in *yku70*Δ (A) and *yku70*Δ *exo1*Δ (D) cells before and after release into cell cycle. (B and E) Southern blotting analysis of the 3’ overhang during once DNA replication in *yku70*Δ (B) and *yku70*Δ *exo1*Δ (E) cells. Magenta arrow: overlapped daughter telomeres with long 3’ overhang, blue arrow: leading telomeres with blunt end, yellow arrow: Exo I trimmed daughter telomeres. (C) Schematic illustration of the E-TRF structures after DNA replication in *yku70*Δ cells. Leading 1, lagging 1 and lagging 2 daughter telomeres refer to the overlapped signals indicated by magenta arrow in (B); leading 2 daughter telomeres refer to the shortened band indicated by blue arrow in (B); the size of each strand (*in vitro* Exo I trimmed, including leading 1, lagging 1 and lagging 2 daughter telomeres with longer overhang) is labeled on the right, and refer to the bands indicated by yellow arrow in (B). See also Figure S7

Further denatured Southern blotting showed that after replication, most of the CA strand was 83 nt (Figure S7B, 120 min), an indication of limited resection to the Rap1 binding site (Figure 1A). The TG strand was measured as 103-107 nt and 83 nt after replication, and the 103-107 nt product converted to 87 nt upon *in vitro* Exo I digestion (Figure S7C, 120 min). These results align with those presented in the native gel of Figure 4B (120 min), strongly support the proposed model, in which three out of the four daughter telomeres contain a ∼24 nt 3’ overhang, and one (daughter leading 2) contains a blunt end (Figure 4C). Of note, the 4 nt length difference of TG strand between daughter leading 2 (83 nt) and other daughter telomeres (∼87 nt after Exo I treatment) (Figure S7C) is likely due to incomplete *in vitro* Exo I digestion of the long overhangs in de-capped daughter telomeres (Figures S2E and S2F).

In the dividing *exo1*Δ (Figures S7D and S7E) and *yku70*Δ *exo1*Δ (Figures 4D and 4E) cells, both the length and structure of telomeres were very different from those in *yku70*Δ cells (Figure 4B), but similar to those in WT cells (Figure 2b). Notably, the 83 bp band observed in WT cells (Figure 2B, 120 min, Exo I+) was not detected in *exo1*Δ and *yku70*Δ *exo1*Δ cells (Figures S7E and 4E), indicating CA strand resection is mediated by Exo1. These results support the conclusion that both the blunt-ended leading strand telomere and the 3’-protruding lagging strand telomeres are shielded by Yku, effectively preventing Exo1-mediated 5’ resection of the CA strand.

### The RNA primer at lagging strand telomeres is removed after replication

For lagging strand telomeres, the RNA primer is presumably removed and expose the 3’ overhang for Cdc13 binding and protection^49^. To address whether an RNA primer is absent on the lagging strand telomere, we examined end structure of telomeres combined with RNase H to digest the DNA-RNA hybrids *in vitro* (Figures S8A and S8B). In WT cells, after replication (Figure 5A), the telomere structures detected without RNase H treatment were similar to these treated with RNase H (Figures 5B and 5C, pink arrow). These results demonstrate that the RNA primer on terminal Okazaki fragment, approximately 10 nt in length, has been removed during or after lagging strand telomere replication, and the RNA primer synthesized by Polα-primase is likely to be initiated at the 3’ end of the TG_1-3_ template strand.

**Figure 5.**
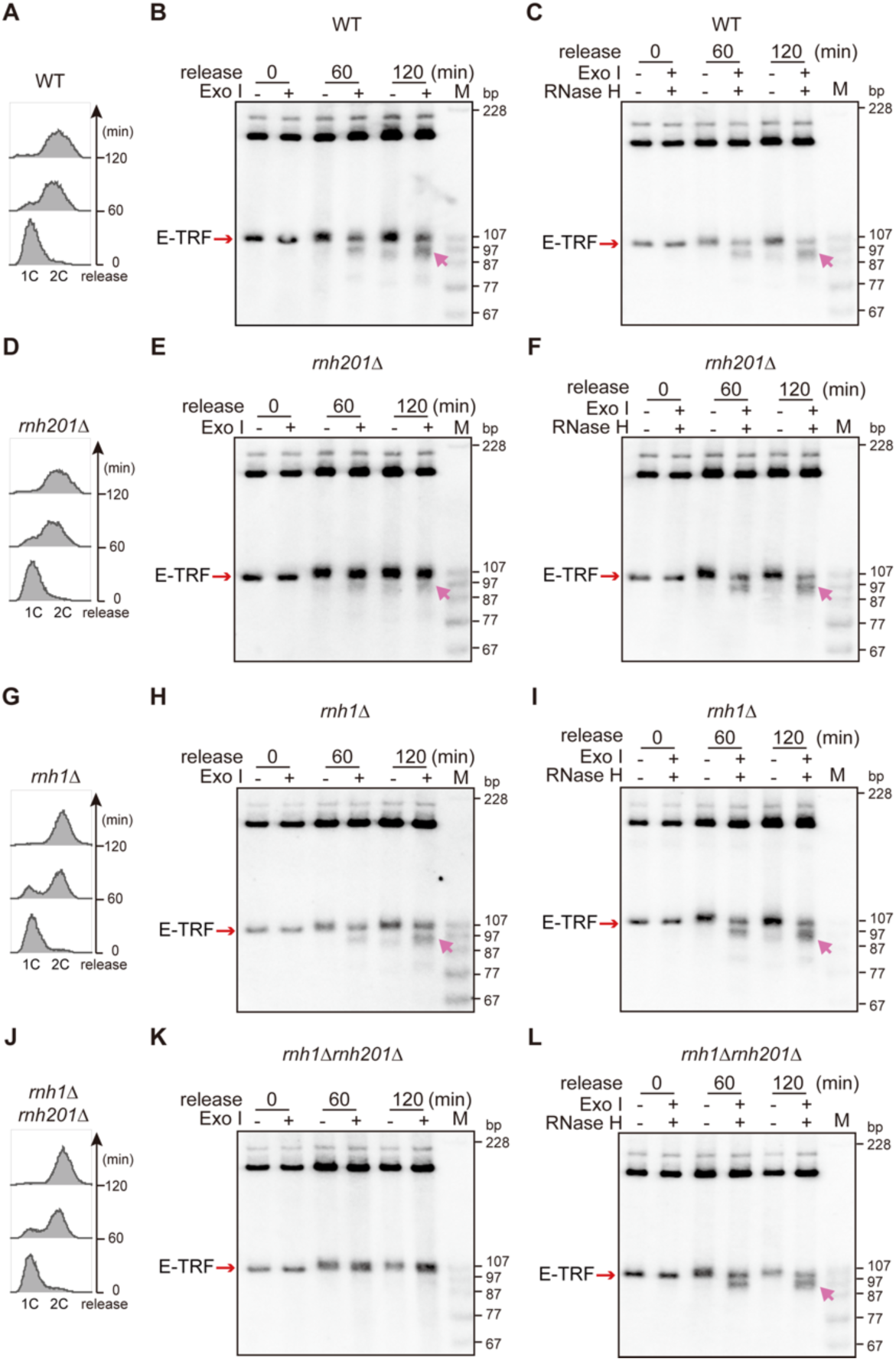
The RNA primer is retained on lagging-strand telomere in *rnh201*Δ cells. (A, D, G and J) FACS analysis of DNA content in WT (A), *rnh201*Δ (D), *rnh1*Δ (G) and *rnh1*Δ *rnh201*Δ (J) cells. (B, E, H and K) Southern blotting analysis of the 3’ overhang at both the leading and lagging strand telomere ends during once DNA replication in WT (B), *rnh201*Δ (E), *rnh1*Δ (H) and *rnh1*Δ *rnh201*Δ (K) cells, pink arrow: Exo I trimmed lagging telomere. (C, F, I and L) Southern blotting analysis of the 3’ overhang at both the leading and lagging strand telomere ends during once DNA replication in WT (C), *rnh201*Δ (F), *rnh1*Δ (I) and *rnh1*Δ *rnh201*Δ (L) cells, pink arrow: Exo I trimmed lagging telomere. Genomic DNA is pretreated with RNase H as labeled. See also Figures S8 and S9.

### RNase H2 is mainly responsible for the RNA primer removal at the lagging strand telomere

To determine which nuclease is responsible for removing the terminal RNA primer, we generated several mutants in which the RNase H coding genes including *RNH201* and *RNH1* (encoding the catalytic subunit of RNase H2 and H1, respectively), which both function in DNA-RNA hybrid removal^21,22^, were singly or doubly deleted, and examined the structures of replicated leading and lagging strand telomeres. In the *rnh201*Δ mutant (Figure 5D), the leading strand telomere was, as expected, resistant to Exo I treatment due to its blunt end structure; the lagging strand telomere, which was sensitive to Exo I treatment in WT cells, became resistant to Exo I treatment (Figure 5E), indicating the absence of 3’ overhang. Significantly, *in vitro* RNase H treatment resulted in approximately half of the 107 bp E-TRFs to become sensitive to Exo I digestion, generating of an E-TRF band of 97 bp in length (Figure 5F, pink arrow). These results demonstrate that in the absence of RNase H2 activity, the RNA primer is retained on the lagging strand telomere (Figure S8B).

In *rnh1*Δ cells, the lagging strand telomere was efficiently trimmed by Exo I *in vitro*, resulting in a 97 bp E-TRF (Figures 5G-5I, pink arrow), suggesting that it contains a ∼10 nt overhang, and little or no RNA primer is retained after replication in the absence of Rnh1. Interestingly, the faint 97 bp E-TRF band observed in the *rnh201*Δ cells (Figure 5E, pink arrow) was completely undetectable upon additional deletion of *RNH1* (Figures 5J-5L), suggesting that Rnh1 plays a redundant but very minor role in RNA primer removal of the lagging strand telomere. Collectively, these results provided the first evidence that RNase H2 is primarily responsible for the last RNA primer removal on lagging strand telomere.

TERRA (Telomeric repeat-containing RNA) is transcribed by RNA polymerase II, using the telomeric C-rich strand as its template^50^. In budding yeast, TERRA levels are extremely low in WT cells^23^. However, deletion of *RNH201* leads to the accumulation of DNA-RNA hybrids at telomeres^24,51^, which could also lead to resistance of TRFs to Exo I digestion (Figure S9A). To eliminate any potential interference of TERRA with telomere structure, we inserted the transcriptional terminator of *CYC1* gene between *KanMX* and the telomeric TG_1-3_/C_1-3_A sequence to block TERRA expression (Figure S9B). The end structures of the replicated telomeres were not affected by the *CYC1*^T^ insertion in either WT or *rnh201*Δ cells (Figures S9C-9H; pink arrow). These results indicate that it is not the TERRA DNA-RNA hybrid, but the DNA-RNA primer hybrid that renders the lagging strand resistant to Exo I digestion. We thus conclude that the ∼10 nt RNA primer was retained on the lagging strand after telomere replication in *rnh201*Δ cells.

### RNA primer removal on lagging strand mainly contributes to end replication problem

To exclude the possibility that terminal 6 bp non-TG sequence affected the processing of the daughter telomeres, we examined length changes of *de novo* telomere in the telomerase-null cells during approximately 25 rounds of consecutive population doublings (PDs). In this process, telomeres gradually shortened and lost the non-TG sequence, and should behave in the same way as native telomeres. If the leading strand usually keeps blunt end, the telomere will shorten at a rate of 2.5 bp per generation (a quarter of the length of the RNA primer), we termed it the “incongruent end model” (Figure S10A). If the leading strand undergoes end processing, the telomere will shorten at a rate of 5 bp per generation (half the length of the RNA primer), which is known as the fill-in model (Figure S1 and S10B). The B-TRFs (BamH I digested terminal fragments) in *tlc1*Δ cells showed signals of scattered ladders, and progressively shortened to between 190 and 235 bp at ∼25 PDs (Figure 6A). Notably, this shortening pattern fits well with our incongruent end model, which shows that telomeres shorten at an average rate of 2.5 bp per generation, and the shortest telomere after 25 PDs is 175 bp (Figures S10A, S10C and S10D), close to the observed 190 bp (Figure 6A). In contrast, the average shortening rate calculated from the fill-in model is 5 bp per generation and the shortest telomere after 25 PDs is 60 bp (Figures S10B-10D), much faster than the experimental data (Figure 6A). Moreover, our results are also consistent with other published data which show that telomeric DNA undergoes a 2 to 3 base pair shortening at each cell division^11,18^. These results suggest that the blunt end leading strand telomeres contribute little to the end replication problem.

**Figure 6.**
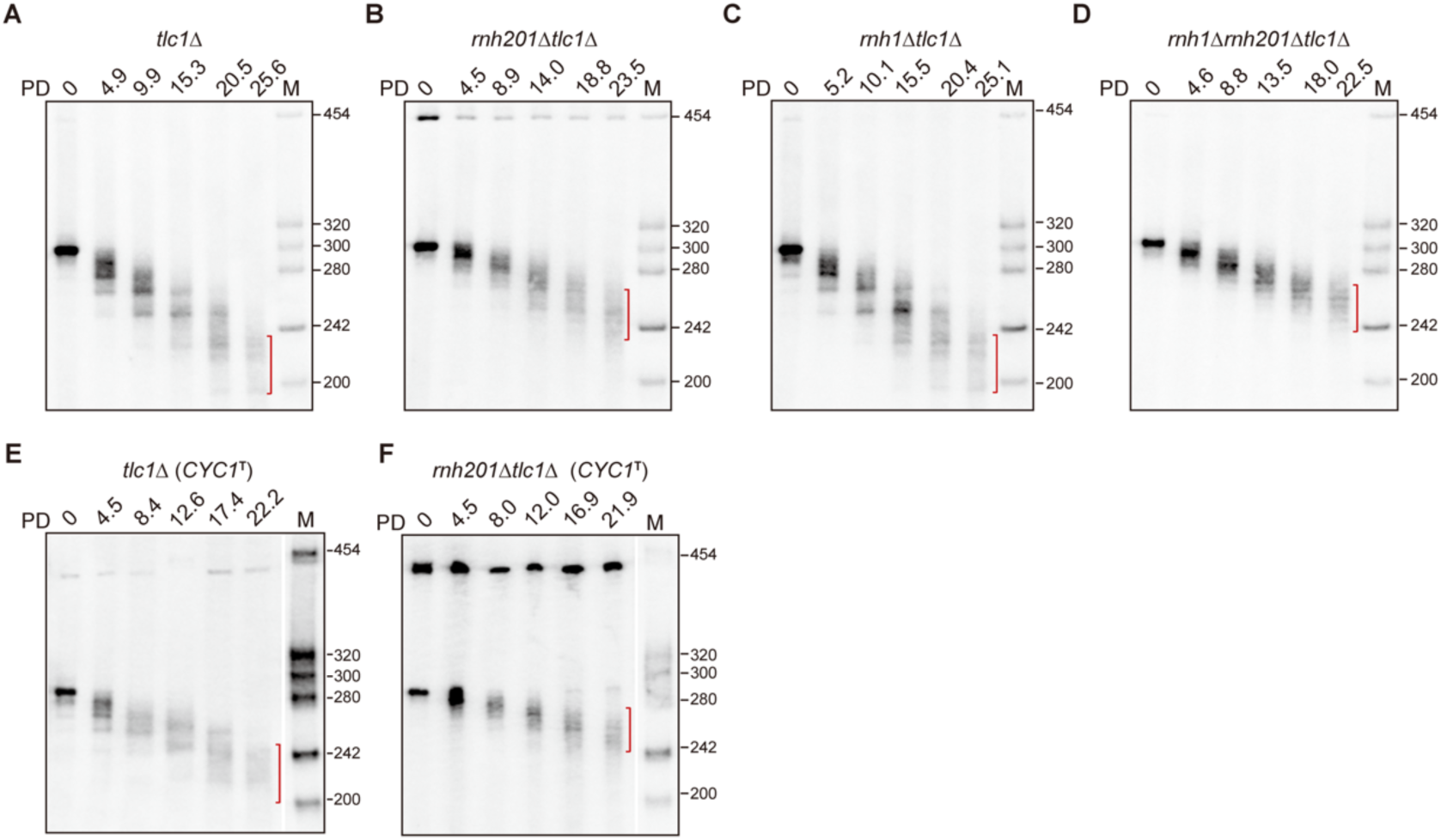
RNA primer retention alleviates telomere shortening independently of TERRA expression. (A-D) Southern blotting analysis of the *de novo* telomere shortening during ∼25 consecutive population doublings (PD) in *tlc1*Δ (A), *rnh201*Δ *tlc1*Δ (B), *rnh1*Δ *tlc1*Δ (C) and *rnh1*Δ *rnh201*Δ *tlc1*Δ (D) cells. DNA is digested with BamH I, *de novo* telomere length is indicated by red bracket. (E and F) Southern blotting analysis of the *de novo* telomere shortening during ∼22 consecutive population doublings (PD) in *tlc1*Δ (*CYC1*^T^) (E) and *rnh201*Δ *tlc1*Δ (*CYC1*^T^) (F) cells in absence of TERRA expression, DNA is digested with BamH I, *de novo* telomere length is indicated by red bracket. See also Figures S10 and S11.

As end replication problem is mainly caused by RNA primer removal, we speculated that the RNA primer retention on lagging strand might prevent telomere shortening during successive cell divisions. To test this hypothesis, we thus examined the shortening of *de novo* telomere in RNase H mutants.

Interestingly, compare to *tlc1*Δ cells, the signals of B-TRFs in the *rnh201*Δ *tlc1*Δ and *rnh1*Δ *rnh201*Δ *tlc1*Δ cells were less dispersed, and gradually shortened to between 230 and 260 bp at ∼25 PDs (Figures 6B and 6D). But the telomere shortening rate in *rnh1*Δ *tlc1*Δ cells is comparable to that in *tlc1*Δ cells (Figures 6A and 6C). Like in telomerase proficient cells, the RNA primer in the *tlc1*Δ and *rnh1*Δ *tlc1*Δ cells was removed (Figures S11A-11C and S11G-11I), but retained on lagging strand in the *rnh201*Δ *tlc1*Δ and *rnh1*Δ *rnh201*Δ *tlc1*Δ cells (Figures S11D-11F and S11J-11L), confirming that RNase H2, but not RNase H1, is mainly responsible for last RNA primer removal in telomerase-null cells. These results indicate that the RNA primer retention counteracts telomere erosion.

Previous studies have shown that TERRA forms R-loops at telomeres to regulate telomere length^51^. It remains possible that the decrease of telomere shortening rate in *rnh201*Δ mutants is due to the TERRA-mediated R-loop. To avoid the interference of TERRA R-loop with telomere length, we inserted the transcriptional terminator of the *CYC1* gene (*CYC1*^T^) next to the *de novo* telomere sequence in *tlc1*Δ and *rnh201*Δ *tlc1*Δ mutants. Elimination of TERRA expression did not change the shortening rate of the *de novo* telomere in telomerase-null *tlc1*Δ and *tlc1*Δ *rnh201*Δ cells (Figures 6E and 6F), compared to no *CYC1*^T^ insertion cells (Figures 6A and 6B), suggesting that RNase H2 inactivation counteracts telomere shortening independently of the TERRA R-loop.

### RNA primer retention at telomeres delays cellular senescence

Based on these observations, we conclude that there are two types of telomere ends in yeast, one with a blunt end and the other with a 3’ overhang (Figure 7A), which are the products of leading and lagging strand replication, respectively. During the telomere replication, if the parental telomere has a blunt end (e.g. a product of leading strand replication), after one round of replication, the leading strand telomere is fully replicated and contains a Yku-shielded blunt end; the lagging strand telomere, whose RNA primer is removed by RNase H2, CA strand shortens 10 nt, generating a 3’ overhang protected by Cdc13 as well as Yku at single-and double-stranded junction (Figure 7B, left panel). If the parental telomere contains a 10 nt overhang (e.g. a product of lagging strand replication) (Figure 7B, right panel), the lagging strand telomere uses the longer TG strand as template, generating a 10 nt 3’ overhang after the RNase H2 mediated RNA primer removal, which is same as parental telomere; however, leading strand telomere uses the 10 nt shorter CA strand as template, its full replication yields a blunt-ended product that is 10 bp shorter than the parental telomere (Figure 7B, right panel). Thus, this incongruent end model emphasizes that the RNA primer removal in the lagging strand replication is the primary cause of telomere shortening, the consequences of which become increasingly severe in the subsequent cycles of leading strand telomere replication.

**Figure 7.**
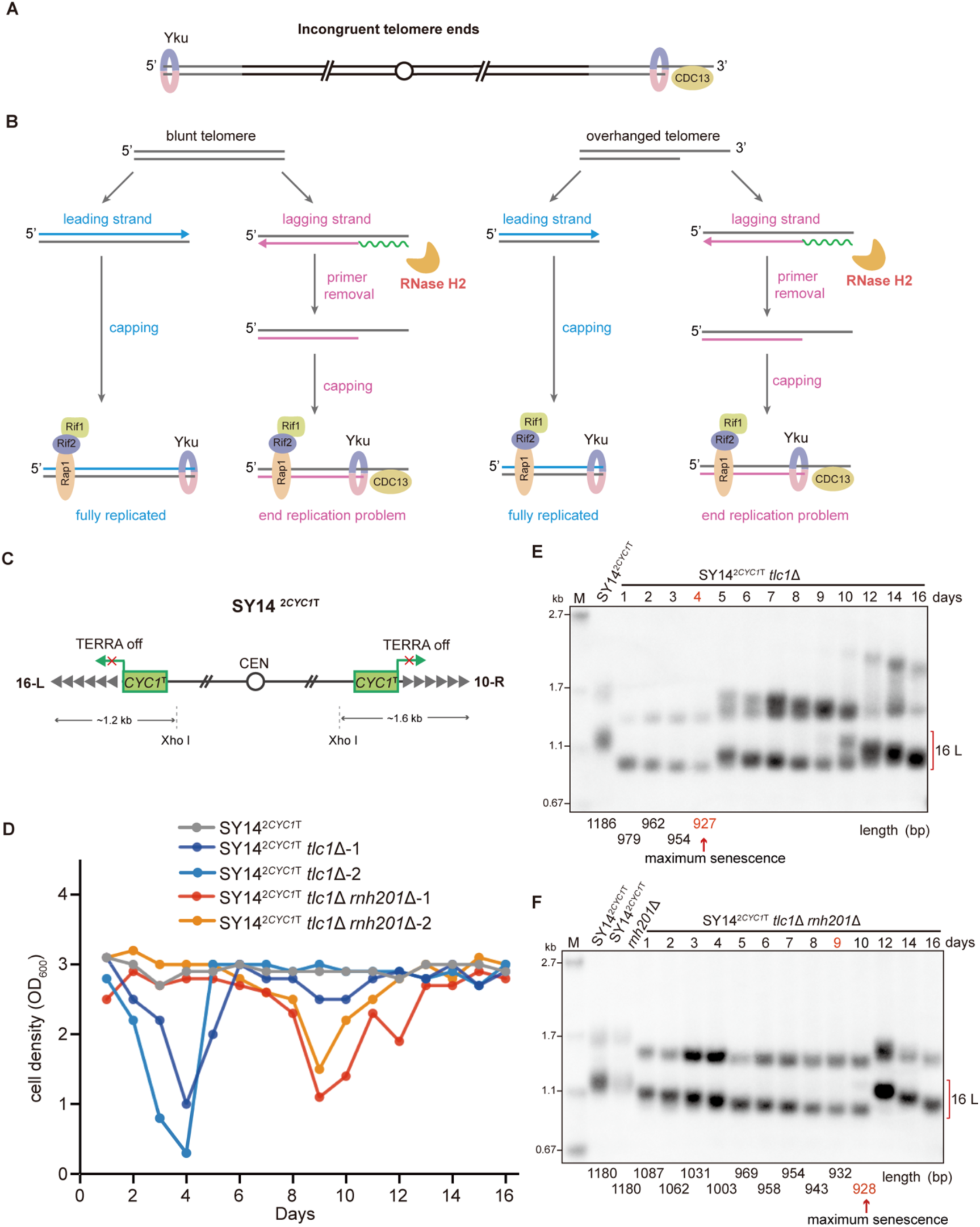
RNA primer retention delays senescence. (A) Schematic of incongruent telomere ends. (B) ‘Incongruent end model’ of telomere replication (see the text for detailed description). (C) Schematic illustration of the insertion of the *CYC1* terminator (*CYC1*^T^) proximal to the two native telomeres (represented by gray triangles) in single chromosome yeast strain SY14^2*CYC1*T^. The terminal fragments digested by Xho I from the left (tel 16 L) and right (tel 10 R) telomeres are approximately 1.2 and 1.6 kb in length, respectively. (D) Cellular senescence analysis of SY14, SY14^2*CYC1*T^ *tlc1*Δ and SY14^2*CYC1*T^ *rnh201*Δ *tlc1*Δ cells during liquid passages. (E and F) Southern blotting analysis of native telomeres in SY14^2*CYC1*T^ *tlc1*Δ (E) and SY14^2*CYC1*T^ *rnh201*Δ *tlc1*Δ (F) cells shown in (D) that underwent senescence and survivor generation. DNA is digested with Xho I, mean length of 16 L telomere TRF analyzed by Image lab is labeled at the bottom, the point of maximum senescence during passage is pointed by red arrow. See also Figures S12 and S13.

In the absence of telomerase, telomeres gradually shorten and cells undergo senescence when telomeres become critically short^12^. As RNA primer retention attenuate telomere shortening (Figures 6A and 6B), would it also delay senescence of telomerase-null cells? We then passaged the *rnh1*Δ *tlc1*Δ and *rnh201*Δ *tlc1*Δ cells on solid plate or in liquid medium, with *tlc1*Δ cells serving as a control. On solid plate, the *rnh201*Δ *tlc1*Δ cells senesced at the 4^th^ restreak, later than the *tlc1*Δ and *rnh1*Δ *tlc1*Δ cells, which senesced at the 3^rd^ restreak (Figure S12A), accounting for a 20-25 generation delay. In liquid medium, the *rnh201*Δ *tlc1*Δ cells senesced at the 6^th^ passage, later than the *tlc1*Δ and *rnh1*Δ *tlc1*Δ cells, which senesced at the 4^th^ passage (Figure S12B), followed by survivor generation^13^. Further analysis of bulk telomeres during liquid passages revealed that the telomere recombination signals in *rnh201*Δ *tlc1*Δ survivors emerged later than those in *tlc1*Δ and *rnh1*Δ *tlc1*Δ survivors (Figure S12C-12E)^13^, consistently suggesting that RNA primer retention at telomeres by RNase H2 inactivation delays telomere-length-dependent senescence.

Previous studies have shown that in absence of both RNase H1 and RNase H2, TERRA DNA-RNA hybrid accumulation delay cellular senescence by promoting homologous recombination^24^. To avoid the interference of the TERRA DNA-RNA hybrids in *rnh201*Δ cells and to optimize the detection of telomeres, we constructed a single-chromosome TERRA-free yeast strain (namely SY14^2*CYC1*T^)^52^. The transcriptional terminator of the *CYC1* gene was inserted immediately upstream of the two remaining native telomeres (tel 16 left and 10 right, 16-L and 10-R) of the single chromosome, respectively, to block the expression of TERRA (Figure 7C). Further single deletion of *TLC1* or double deletion of *TLC1* and *RNH201* resulted in senescence when SY14^2*CYC1*T^ *tlc1*Δ and SY14^2*CYC1*T^ *tlc1*Δ *rnh201*Δ mutants were inoculated in liquid medium at passage 4 and 9, respectively (Figure 7D), indicating that *RNH201* deletion significantly delayed senescence by ∼50 generations. Telomere length analysis showed that TRFs of the tel 16-L telomere of SY14^2*CYC1*T^ *tlc1*Δ cells shortened abruptly by 207 bp at the first passage (from 1186 bp to 979 bp), then shortened slowly to a critical point (927 bp) at the passage 4, and telomeric TG_1-3_ recombination took place at the passage 5 (Figure 7E), consistent with survivor generation. In contrast, TRFs of the tel 16-L telomere of SY14^2*CYC1*T^ *tlc1*Δ *rnh201*Δ cells shortened by only 93 bp during the first passage (from 1180 bp to 1087 bp), and took 8 passages to shorten to a critical point (928 bp), and finally underwent recombination until the passage 10 (Figure 7F). These results indicate that Rnh201 inactivation attenuates telomere shortening and delays cellular senescence independently of the TERRA DNA-RNA hybrid.

## DISCUSSION

Telomeres, situated at the ends of chromosomes, exhibit characteristics resembling double-strand breaks, making them susceptible to resection activities. However, telomeres also possess unique and relatively stable structures. A telomere in *S. cereviaise* is thought to consist of a Cdc13-protected G-rich 3’ overhang, which is presumably resulted from removal of a hypothetical terminal RNA primer and/or limited 5’ C-rich strand resection^3,4^. This cursory understanding sounds too intuitive to refute, given that the variation in telomere length between cells and the ultrashort nature of the RNA primer make it difficult to precisely monitor telomere end processing. The *de novo* telomere system, initially developed by Gottshling’s group, offers an appealing approach for dissecting telomere end processing events at a particular telomere^31^. In addition, short *de novo* telomeres are preferential targets of telomerase, but long *de novo* telomeres featuring extended TG_1-3_/C_1-3_A sequences (>200 bp) circumvent the interference of telomerase and homologous recombination activities, both of which typically favor shorter telomeres^17,30,53,54^. Moreover, the engineered long *de novo* telomeres induced by CRISPR-Cas9 in this work closely resemble natural telomeres, making them ideal for studying telomere replication and processing.

It was somewhat unexpected to see a stable blunt end at the *de novo* telomere in the G2/M and G1 phase cells (Figure 1). It was truly surprising to see both a stable blunt end of the leading strand telomere after replication (Figure 2) and the prevalence of blunt ends on native telomeres (Figure 3). These observations challenged the widely accepted fill-in model that consistently depicts protruding 3’ overhangs at both leading and lagging strand telomeres (Figure S1)^4,10,55,56^. There could be several reasons why the blunt-end telomere was not determined for a long time. (1) Telomerase-mediated telomere elongation typically requires a 3’ overhang, reinforcing the expectation that a telomere should be structured to facilitate telomerase action^1,14,57^ (2) Telomeres of different lengths may be subject to different end processing^4,30,58^. In particular, when a telomeric TG strand is elongated by telomerase, the subsequent synthesis of the CA strand by conventional DNA polymerases (such as Polα/primase and Polδ) usually leaves a 3’ overhang after RNA primer removal^14,16,59^. Thus, previously used assays likely mixed up the signals from telomerase elongated telomeres and telomeres replicated by conventional semi-conservative DNA replication in a cell cycle. (3) The widely used high resolution primer-ligation extension assay and single telomere length analysis (STELA)^3,56^ were based on the assumption of a 3’ overhang, failed to detect telomeres with a blunt end, and unable to distinguish between the leading strand and lagging strand. (4) Due to the heterogeneity of telomere length, the rough measurement of 3’ overhangs on bulk native telomeres (e.g. by the TPX, telomere PCR/Exo I digestion) assay is too coarse to verify the length before and after Exo I digestion of a single telomere^4^, the judgement of the average length is not accurate as telomeres of different lengths are faced with completely different end processing^4,30,58^.

The *de novo* telomere formation system used in this study is capable of generating a long telomere of designed homogeneous length (250 bp, Figure 1), which not only avoids the drawbacks of previously established methods, but also allows us to directly detect the length of 3’ overhangs on both the leading and lagging strand telomeres simultaneously at near-nucleotide resolution (Figures 2 and 4), and to identify mutants with defects in RNA primer removal (Figure 5). In addition, the high-resolution genome-wide native END-seq ensures unbiased detection of blunt-ended telomeres *in vivo*, and provides additional evidence to support the conclusion that leading and lagging strand telomeres have incongruent structures (Figure 3).

Blunt telomeres appear to be optimal targets for Yku, which plays a role in inhibiting chromosome end resection^36,60^. In line with this, the absence of Yku renders the leading strand telomere susceptible to 5’ resection (Figures 4B and 4C). Although previous studies had shown that blunt-ended leading strand telomeres exist in plants and mammals^61,62^. It remains to explore whether the protection and blunt end structure of leading strand telomeres observed in *S. cerevisiae* will hold true in other organisms^63–66^.

During genome duplication, replication of the lagging strand is discontinuous^67^. Our results found that the 8 to 12 nt RNA primer initiated at the end of the parental lagging strand telomere is primarily removed by RNase H2 (Figures 5E and 5F). The identification of Rnh201 mediated terminal RNA primer removal aligns with previous findings that Rnh201 degrades the RNA moiety of DNA-RNA hybrids on telomeres^51^ and involved in Okazaki fragment maturation during DNA replication^26,27,68^. Nevertheless, it remains possible that other nucleases (such as Rnh1) are redundantly involved in RNA primer removal at telomeres (Figures 5K and 5L). The length of the RNA primer is consistent with the generally accepted model^4,10,55^, and 10 nt 3’ overhang falls within the substrate preference for Cdc13^34,49^. Yku also protects the lagging strand telomere from Exo1-mediated extensive resection (Figures 4B-4E), supporting the notion that Yku binds at the junction of dsDNA and ssDNA of a telomere and inhibits 5’ resection^36,69,70^. It is noteworthy that the position of the RNA primer differs in mammals, as the RNA primer does not initiate immediately from the end in both the semi-conservative DNA replication and telomerase elongation pathways in mammals^14,59,71,72^. Furthermore, although the precise efficiency of RNA primer removal is difficult to assess, the relative ratio of overhanged lagging strand and blunt-ended leading strand after replication is approximately 1:1, indicating efficient removal of the majority of RNA primers. Deletion of either *RNH201* or *RNH1* or both does not change telomere length in telomerase-proficient cells (Figure S13A)^13^,and the telomere length in telomerase hyperactive *rif2*Δ and *rif2*Δ *rnh201*Δ cells is similar (Figure S13B)^73,74^, suggesting that RNA primer retention has little effect on telomerase activity.

In absence of RNase H2, the delay of cellular senescence was suggested to be attributed to the accumulation of TERRA DNA-RNA hybrids which promote HR-mediated telomere elongation in previous studies^24,25,51^. However, although the RNase H1 and RNase H2 both function in the removal of TERRA DNA-RNA hybrids^24^, either slowed telomere shortening or delayed senescence was only observed in *rnh201*Δ *tlc1*Δ cells, but not in *rnh1*Δ *tlc1*Δ cells (Figures 6B, 6C and S12), indicating that RNase H2 and RNase H1 regulates telomere homeostasis differently. Consistently, elimination of TERRA expression in telomerase-proficient or-deficient *rnh201*Δ cells has no epistatic effect on telomeric RNA primer retention, telomere shortening rate or cellular senescence progression (Figures S9, 6E, 6F and 7C-7F). Our results elucidate the mechanism of last RNA primer removal by RNase H2. The retention of RNA primer at the telomeres significantly attenuates telomere shortening and delays senescence in the absence of *RNH201* (Figures 6 and 7).

It has been widely accepted that the leading strand telomere also contains a 3’ overhang^3,9^. This overhang was thought to result from a process involving 5’ resection of the blunt intermediate of replication, fill-in synthesis, and RNA primer removal (Figure S1)^4,45,75^. As a result, the end replication problem, which occurs at both chromosome ends, is at least twice as severe as at one end (Figures S10A and 10B). However, our results show that the fully replicated leading strand telomere terminates with a Yku-capped stable blunt end (Figures 2 and 4), perfectly recapitulating the original proposal that the lagging strand telomere is the primary culprit behind the end replication problem due to the intrinsic removal of the RNA primer (Figure 7)^76^. Consistently, observations from our study and other published data suggest that telomeric DNA undergoes a 2 to 3 base pair shortening at each cell division (Figures 6A and S10A)^11,18^, approximately a quarter of the length of an RNA primer, which had been also detected in some human cells^61^. Thus, we conclude that the fully replicated leading strand telomere is protected by Yku and remains relatively stable, making a minor contribution to the end replication problem.

When the RNA primer is retained at the telomeres, as seen in *rnh201*Δ cells, the rate of telomere shortening is further reduced (Figure 7). An intuitive explanation for the slowed telomere shortening is that although the RNA primer retention on the lagging strand impedes telomere shortening within a single cell cycle, the RNA primer situated at the extreme terminus of the CA strand cannot function as a template for replication in subsequent cell cycles. This limitation arises from the DNA polymerase’s inherent preference against utilizing RNA as a template, ultimately leading to telomere shortening over time. But the 5’ end of the DNA-RNA primer hybrid, which is refractory to exonuclease activities (e. g. Mre11 or Exo1)^77^, plays a protective role to attenuate telomere shortening.

### Limitations of the study

Although we have proposed an incongruent-end model of telomere replication in budding yeast, the length of telomeres and the binding proteins and end processing in yeast are quite different from the telomeres in human cells. Whether the blunt end leading strand also exists in human cells remains to be determined. In addition, the 3’ overhang is much longer in mammals than that in yeast, and it is unclear whether the mechanism of the last RNA primer removal is the same in mammals as in yeast. Moreover, telomerase in yeast could act on a telomere to elongate the G-strand, followed by C-strand synthesis by Polα-primase and Polδ. Whether the RNA primer on a telomerase-elongated telomere is also removed by RNase H2 remains to be clarified.

## RESOURCE AVAILABILITY

### Lead contact

Further information and requests for strains should be directed to Jin-Qiu Zhou

### Materials availability

All yeast strains and DNA constructs generated in this study are available upon request.

### Data and code availability

Raw and processed END-seq data have been deposited at GEO at accession number GSE283459. Genome sequencing data have been submitted to NCBI database with an accession number of: SRR31637145, assembled genome sequence can be viewed and downloaded through http://www.bioinformaticspa.com/DDA/new_subTelomere_withN.fa.

## ACKNOWLEDGMENTS

We thank Dr. Fei-Long Meng and lab members for helpful discussion. This work was support by grants from Chinese Academy of Sciences (Strategic Priority Research Program, No. XDB0570000) and National Natural Science Foundation of China (No. 32150004) to J.-Q.Z.

## AUTHOR CONTRIBUTIONS

J.-Q.Z. and T.Y. conceived the study, analyzed the data and wrote the manuscript. T.Y. performed the most of the experiments. S.W. and W.W. establish the native END-seq assay, W.W. and Q.Y. analyzed the sequencing data. J.-T.Z. performed the senescence analysis. J.-C.L. and Z.-J.W. constructed the plasmids and discussed the manuscript. J.-Q.Z. supervised the project.

## MATERIALS AND METHODS

### Yeast strains

All yeast strains are listed in Table S1, cells of *de novo* telomere system were derivatives of a *Saccharomyces cerevisiae* yeast strain: ygk171: *MAT*a, his3-1, ura3-1, leu2-3, trp1-1, ade2-1, can1-100^78^. Cas9 expression cassette A *LEU2*-Cas9-gRNA expression cassette was inserted to the *leu2-3* locus; an *ADE2* marker was inserted to the *MNT2* gene locus; a *Kan*MX-TG250 fragment was inserted to the *ADH4* gene locus. Gene knock-in was performed by a PCR-amplified DNA product mediated homologous recombination followed by Lithium acetate transformation, positive clones were verified by PCR^79^, and the resulting strain was hereafter served as WT control strain. Gene deletion mutant(s) was generated by a plasmid (yeast integrating plasmids pRS300 series)^80^ or PCR fragments mediated gene knockout.

### Plasmids

#### Integrating Cas9 plasmid

The TEFp-NLS-Cas9-NLS-CYC1t was PCR amplified from the pCas9 plasmid^52^, the selection marker *URA3* was PCR amplified from the pRS316 plasmid, *URA3*-Cas9 fragment was obtained by overlap PCR, and were integrated into the *leu2-3* locus for the constitutive expression of Cas9. Gal10 promoter was PCR amplified from the pCas9 plasmid^52^, gRNA-CYC1t was PCR amplified from the pCgRNA plasmid^52^, gRNA sequence (5’-ATATTATCCCATTCCATGCG-3’, PAM: TGG) was introduced by overlap PCR, the selection marker *LEU2* was PCR amplified from the pRS315 plasmid. *LEU2-*Gal10p-gRNA-CYC1t fragment was obtained by overlap PCR, and were integrated into the *leu2-3-URA3* locus for the galactose inducible expression of gRNA and replacement of *URA3* marker.

#### Integrating gRNA plasmid

To construct the TG250 integrating plasmid, 500 bp upstream homology arm, 450 bp downstream homology arm of *ADH4* locus and the selective marker *KanMX* were PCR amplified and integrated to a pRS313 plasmid by infusion cloning. The synthesized TG250-gRNA cassette was released from the cloning vector by NotI and EcoRI digestion, and then ligated to the pRS313-*ADH4*-*KanMX-ADH4* plasmid. *ADH4*-*KanMX-*TG250-gRNA*-ADH4* fragment was obtained by PCR for yeast transformation, positive clones were verified by PCR identification, Southern blotting and Sanger sequencing.

#### *De novo* telomere induction and cell cycle monitoring

Cells were cultured in synthetic media without adenine (2% glucose). For *de novo* telomere formation assay, prior to induction, cells were diluted in YEP-2% raffinose for 4 hours to reach log phase, then cells were synchronized for 2 hours by adding nocodazole to a final concentration of 5 μg/ml, or α-factor to a final concentration of 50 ng/ml for 2 hours, respectively^81^. Finally, galactose was added to the medium to induce gRNA expression^31^, and cells were harvested at the indicated time. Cell cycle arrest was confirmed by FACS analysis. For the cell cycle release assay, cells were first cultured with galactose for 3 hours to induce gRNA expression and *de novo* telomere formation. They were then arrested in early S phase with hydroxyurea (HU, a final concentration of 250 mM) for 2 hours. Glucose was added to completely block gRNA expression for 2 hours. HU was then washed out, and cells were released into the cell cycle with fresh YPD medium. Nocodazole was added after 60 minutes to prevent a second round of the cell cycle, and cells were harvested at different time points. Cell cycle progression was confirmed by FACS analysis. For the cell doubling assay, asynchronized telomerase null cells were cultured in galactose for 8 hours, then cells were diluted in YPD medium to ∼OD_600_=0.25, hereafter diluted and collected every 12 hours (∼5 PDs) until 60 hours. Cell doubling was confirmed by measuring OD_600_ using a spectrophotometer^4^.

#### Cellular senescence assay

Single clones from the isogenic *TLC1* deletion strains were used for cellular senescence analysis^82^. Cells were cultured in YPD medium and grown to stationary phase, and cell density (OD_600_) was measured every 24 hours using a spectrophotometer. Cultures were diluted with fresh YPD medium to ∼OD_600_=0.01. Passages were repeated for 14 days (16 days for SY14^2*CYC1*T^cells).

## FACS

Cells were first washed once with sterile ddH_2_O, then thoroughly mixed with 70 % ethanol for fixation, and stored at 4_°_C overnight. Fixed cells were washed with 50 mM sodium citrate (pH 7.2), resuspended in sodium citrate with RNase A (final concentration of 200 μg/ml), and incubated at 37_°_C for 3 hours; Protease K (final concentration of 200 μg/ml) was added, and incubated at 50_°_C for 1 hour. Cells were washed, followed by sonication and stained with PI (propidium iodide, 10 μg/ml) for 1 hour at room temperature. FACS analysis was performed using BD Forteasa^83^.

### Southern blotting analysis

Cells collected from each time points were stored at-80°C, and their genomic DNA was extracted in parallel using a mild protocol^84^. Cells were first digested with zymolyase at a final concentration of 1.5mg/ml in sorbitol (1M) buffer, then were resuspended in lysis buffer, followed by phenol/chloroform extraction and ethanol precipitation. After dissolved in water, DNA was further purified by incubation with RNase A. RNase A treatment was omitted and RNase H (New England Biolabs) digestion was performed at 37°C for 2 hours in the RNA primer detection assay. Extracted DNA was digested with EcoR V or BamH I at 37°C for 2 hours, then Exo I (New England Biolabs) was added, and incubated at 37°C for 1 hour to remove 3’ overhang. The DNA was separated on native PAGE (8% for E-TRF, 6% for B-TRF) or denatured 6% urea (7 M)-PAGE, and Southern blotting was performed as previously described^39^. Bulk telomere analysis is performed as mentioned^82^, briefly, the Xho I digested DNA is separated on 1% agarose gel and transferred to nylon membrane, then hybridized with CA probe. The 3’-biotin-labelled probes contained the sequences 5’-CAC CAC ACC CAC ACA CCA CAC CCA CA-3’-biotin (CA probe) and 5’-TGT GGG TGT GGT GTG TGG GTG TGG TG-3’-biotin (TG probe), respectively. Sequences of the size markers for denatured PAGE are listed in Table S2. All results were repeated three times independently.

### Double-stranded oligonucleotide/markers preparation

The oligonucleotides used in this study were synthesized by the Azenta company, sequences of the oligonucleotides are listed in Table S3. Overhanged oligos were prepared by one-step elongation PCR. Sequences of oligonucleotides and primers are listed in Extended table S3. PCR products were separated on a native PAGE, and extracted from polyacrylamide gel using QIAEX II kit.

### Genome sequencing and assembly

High quality genomic DNA was extracted from WT cells according to the manufacturing instruction. 2 μg genomic DNA was used for the 15 kb SMRT-bell sequencing library construction. The g-TUBE was used to shear gDNA to 6 kb to 20 kb. After shearing, AMPure PB Beads were used to concentrate sheared gDNA. Then followed by single-strand overhang removal, DNA damage repair, end repair and SMRTbell hairpin adapter ligation according to the manufacturing instruction. Nuclease treatment was performed to remove imperfect SMRTbell templates. Finally, followed by size selection with the BluePippin system to remove small insert. The SMRT-bell yeast genomic library was sequenced using PacBio Sequel IIe SMRT Cell 8M with 30-h movie, yielding about 10 Gb of HiFi reads. The HiFi reads were assembled with hifisam(v0.19.9-r616)(PMID: 33526886) and then polished 3 times with minimap2(v2.28-r1209)(PMID: 29750242) and Racon(v1.5.0)(PMID: 28100585). Finally, 16 contigs were assembled using RagTag(v2.1.0)(PMID: 36522651) with S288C genome as a reference.

### END-seq sequencing and analysis

END-seq was performed as previously described^47^. Briefly, cells were synchronized to G2/M phase before harvest, cell cycle arrest was confirmed by cell morphology. ∼10^8^ cells were washed twice by 50 mM EDTA (pH 8.0) and embedded in 0.75% low melting agarose plugs with Lyticase (1 mg/ml, final concentration). Plugs were then treated with Lyticase and Lysing (1 mg/ml, final concentration, 37°C for 1.5 hours), Proteinase K (50°C for 2 hours, followed by 37°C for 7 hours) and RNase A (37°C for 1 hour) in turn. DNA ends within plugs were either not trimmed (native END-seq) or trimmed (typical END-seq) with Exonuclease T and Exonuclease VII. All samples were followed by A-tailing with Klenow fragment exo^-^ to ligate with 3’ T overhang of biotinylated hairpin adaptors (introducing Illumina p5 primers). After plug melting and sonication, DNA fragments were sheared to 300-500 bp in length and captured by streptavidin beads. End-repair by T4 DNA polymerase, Klenow fragment and T4 polynucleotide kinase was performed before second A-tailing with Klenow fragment exo^-^, allowing the ligation of distal hairpin adaptors with p7 Illumina primers. Finally, after USER enzyme digestion, libraries were PCR amplified and recovered for Illumina sequencing. END-seq sequencing reads were firstly trimmed to remove adapters by cutadapt (v4.1) (DOI:10.14806/ej.17.1.200) with parameters-e 0.1-O 3-m 55--quality-cutoff 25, then reads were aligned to the assembled yeast genome using bowtie (v1.2.1.1) (PMID: 19261174) with parameters-n 3-l 50-k 1. To circumvent the interference of internal telomeric sequence, telomeric repeats in sub-telomeric regions are masked with “N”. functions’view’ and’sort’ of samtools (v.1.9) (PMID: 19505943) were used to convert and sort the aligned.sam files to sorted.bam files. The forward reads were retained using samtools’view’ function for subsequent analysis. bam files were further converted to.bed files using the bedtools (v2.25.0) bamToBed command (PMID: 20110278) and bedGraph files were generated using bedtools genomecov, normalized by reads per million (RPM) and then converted into.bigWig files using bedGraphToBigWig from UCSC utilities (PMID: 20639541)

## Supplemental Figures and figure Legends

**Figure S1.**
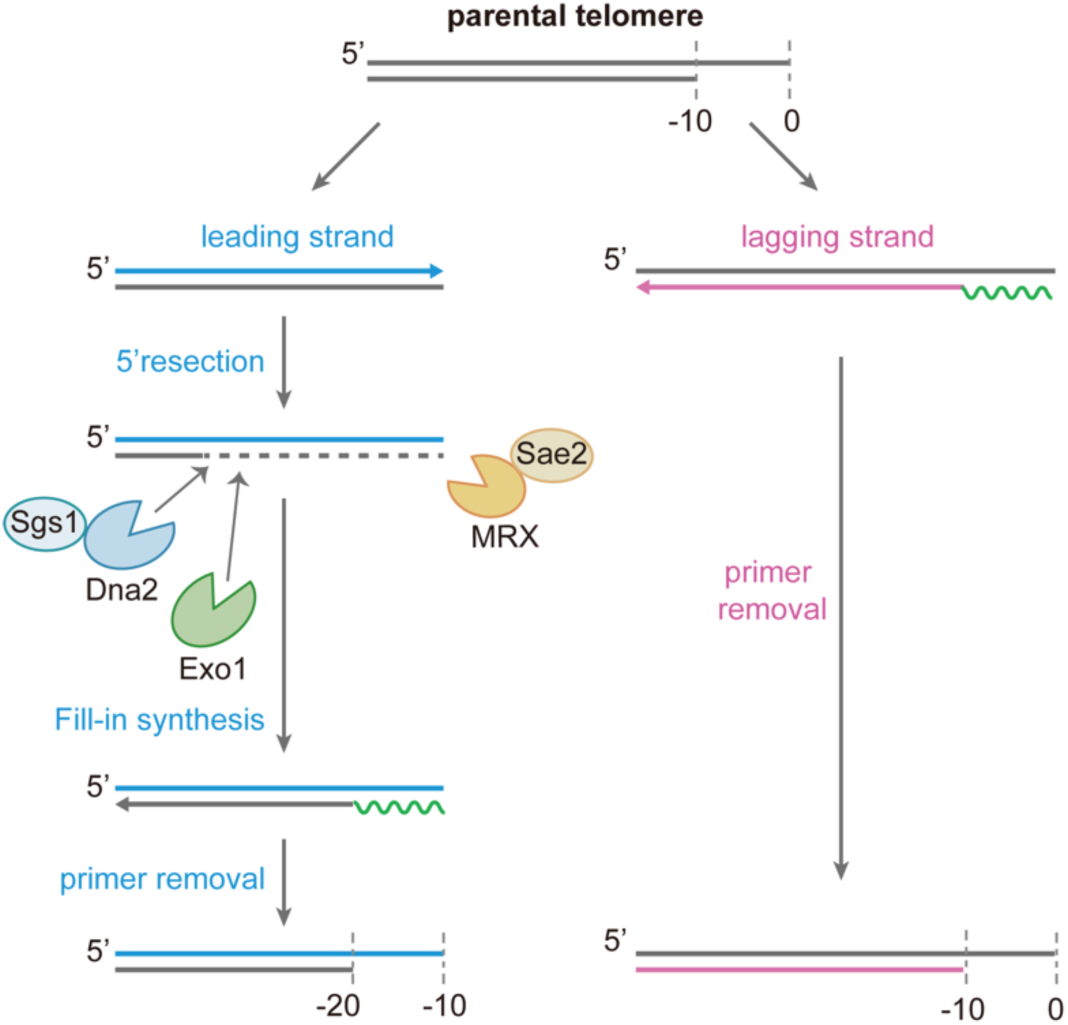
Schematic illustration of the fill-in model of telomere replication. Parental telomeres have a 3’ overhang with hypothetical length of 10 nt. After replication, the RNA primer on the lagging strand telomere is removed, resulting in a protruding 3’ overhang; the leading strand telomere with a blunt end undergoes resection by nuclease such as Mre11, Exo1 and Dna2, followed by fill-in synthesis and RNA primer removal. Finally, both the leading and lagging strands possess 3’ overhangs of approximately 10 nt^1,2^.

**Figure S2.**
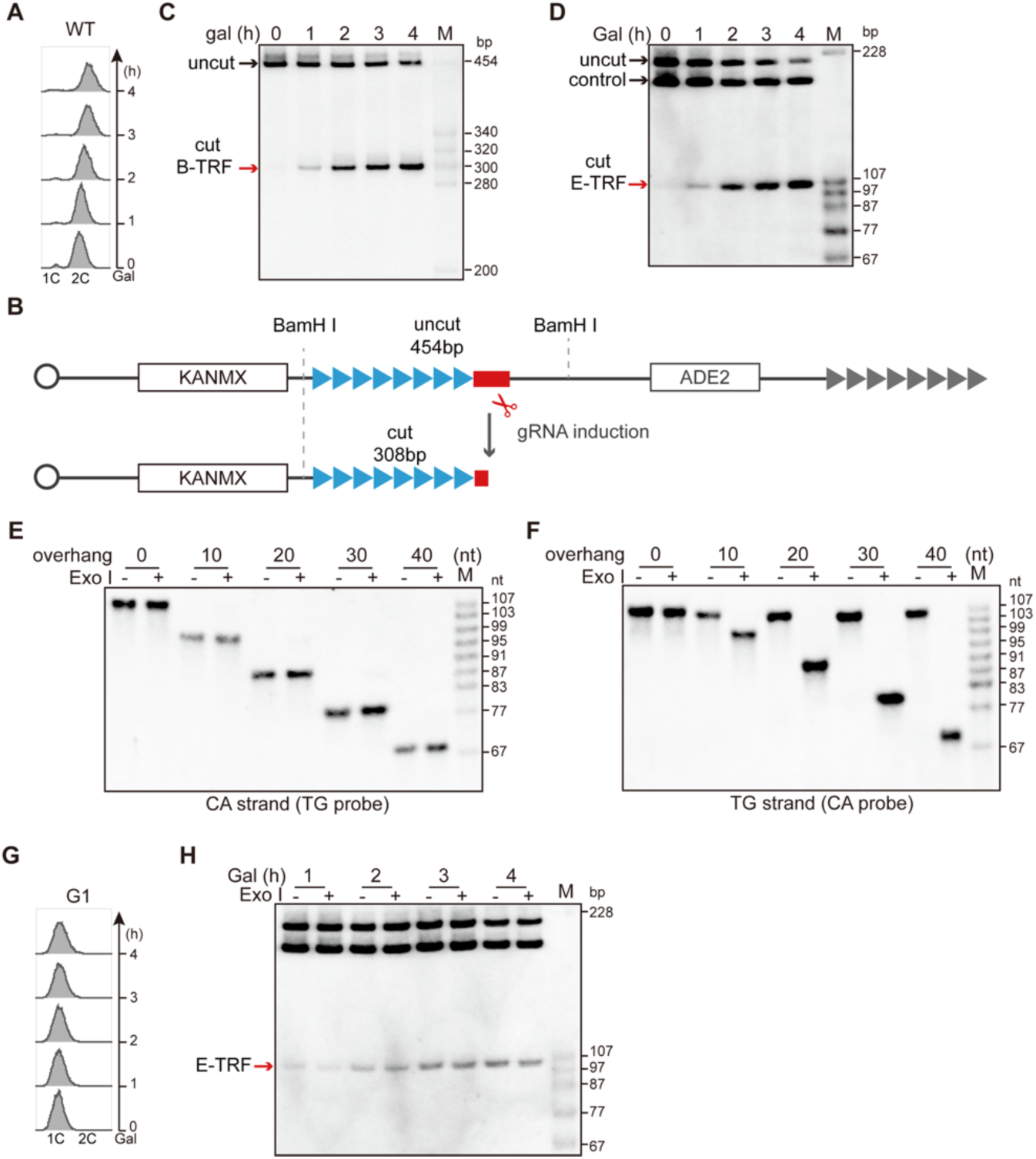
Examination of *de novo* telomere formation and 3’ overhang, related to. Figure 1. (A and G) FACS analysis of DNA content in nocodazole (A) and α-factor (G) arrested WT cells. (B) Schematic illustration of the *de novo* telomere induction system, BamH I sites are displayed. (C) Detection of the *de novo* telomere formation at G2/M phase in WT cells. The genomic DNA is digested with BamH I, Southern blotting was performed by 6% native PAGE, hybridized with telomeric specific CA probe, B-TRF includes 250 bp telomeric DNA of *de novo* telomere and the adjacent non-TG sequence. (D) Detection of the *de novo* telomere end at G2/M phase in WT cells. The genomic DNA is digested with EcoR V, Southern blotting was performed by 8% native PAGE, E-TRF includes 107 bp of *de novo* telomere end. (E and F) Denatured Southern blotting verifying the length of the single-stranded CA strand (E) and TG strand (F) separated on 6% urea-PAGE, hybridized with TG (E) and CA (F) probe, respectively. In line with Exo I activity (3’ to 5’ exonuclease), the single-stranded TG-rich overhangs were trimmed by Exo I (F, Exo I+), but measured a few nucleotides longer than their complementary CA-rich oligos (E, Exo I+), because short 3’ overhangs are not favorable substrates for Exo I. The sizes of the markers are indicated. (H) 3’ overhang detection of *de novo* telomere induced in G1 phase.

**Figure S3.**
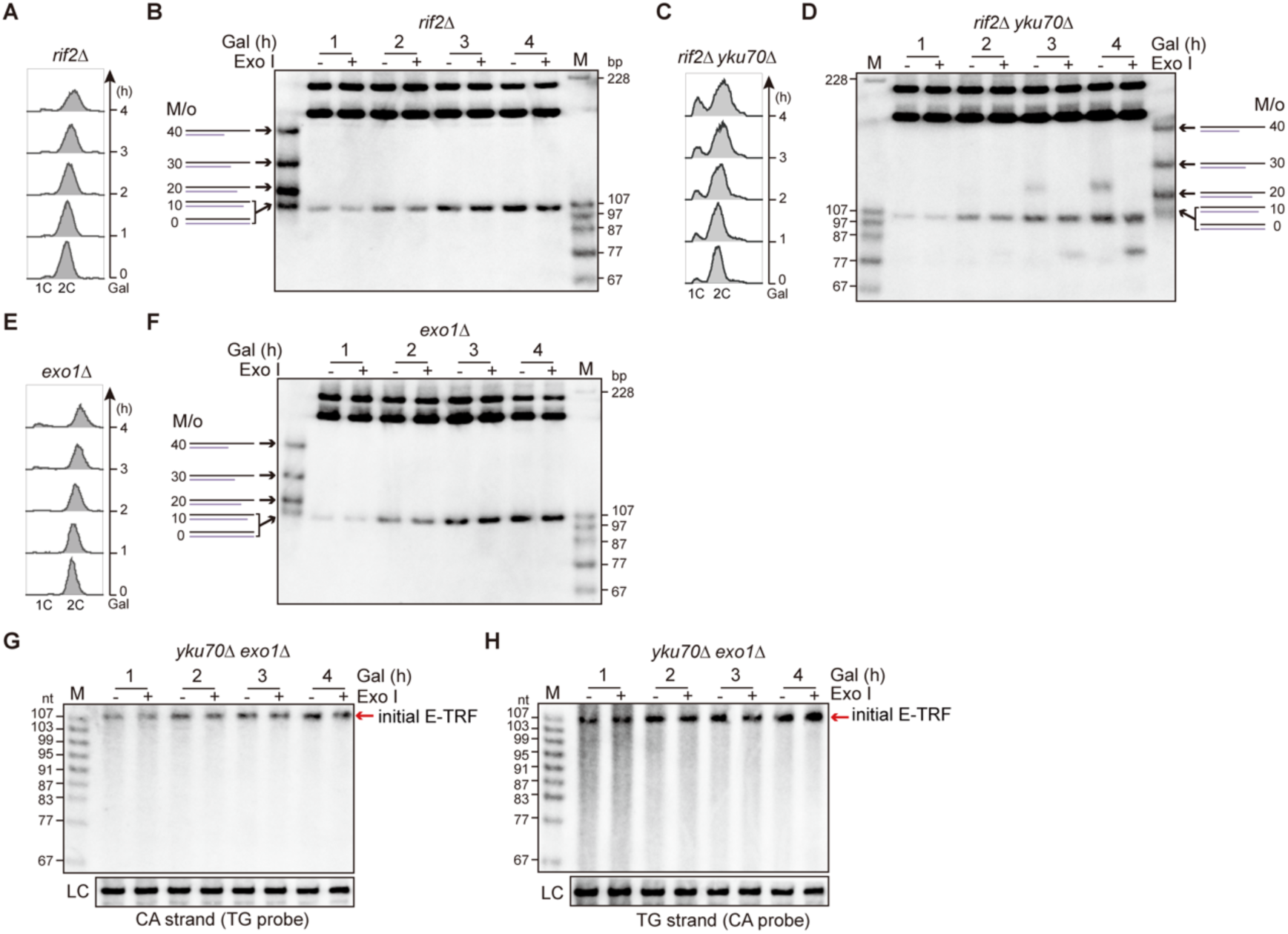
*De novo* telomere is resected by Exo1 in *yku70*Δ cells, related to. Figure 1. (A, C and E) FACS analysis of DNA content of nocodazole-synchronized *rif2*Δ (A), *rif2*Δ *yku70*Δ (C) and *exo1*Δ (E) cells. (B, D and F) Southern blotting analysis of the 3’ overhang at *de novo* telomere in *rif2*Δ (B), *rif2*Δ *yku70*Δ (D) and *exo1*Δ (F) cells. (G and H) Denatured Southern blotting analysis of the CA-strand (G) and TG-strand (H) in *yku70*Δ *exo1*Δ cells at G2/M phase, hybridized with TG probe (G) and CA probe (H), respectively.

**Figure S4.**
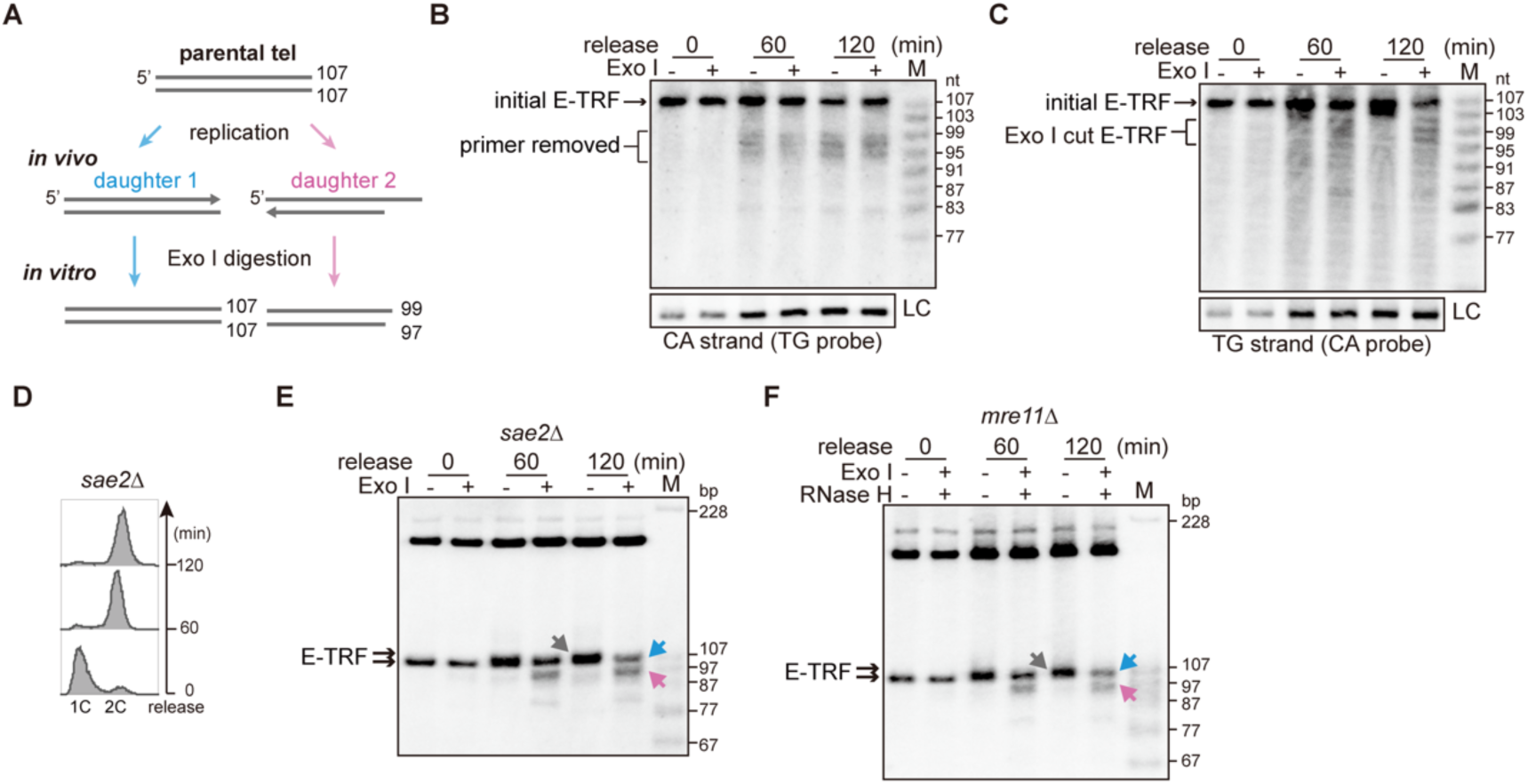
Leading strand telomere is blunt-ended, related to. Figure 2. (A) Schematic illustration of daughter telomeres after the completion of telomere replication. Daughter telomere 1 maintains a stable blunt end, while the daughter telomere 2 has a 3’ overhang of approximately 10 nt. (B and C) Denatured Southern blotting analysis of the CA-strand (B) and TG-strand (C) during once replication in WT cells, hybridized with TG (B) and CA (C) probe, respectively. CA strand indicated by black bracket is the replication product after RNA primer removal (B). TG strand indicated by black bracket is the overhanged telomere, sensitive to Exo I digestion (C). (D) FACS analysis of DNA content in *sae2*Δ cells before and after release in to cell cycle. (E) Southern blotting analysis of the 3’ overhang at both telomere ends during once DNA replication in *sae2*Δ cells. Grey arrow: overlapped daughter telomeres, blue arrow: leading telomere, pink arrow: lagging telomere. (F) Southern blotting analysis of the 3’ overhang at both telomere ends during once DNA replication *mre11*Δ cells, genomic DNA is pretreated with RNase H as labeled.

**Figure S5.**
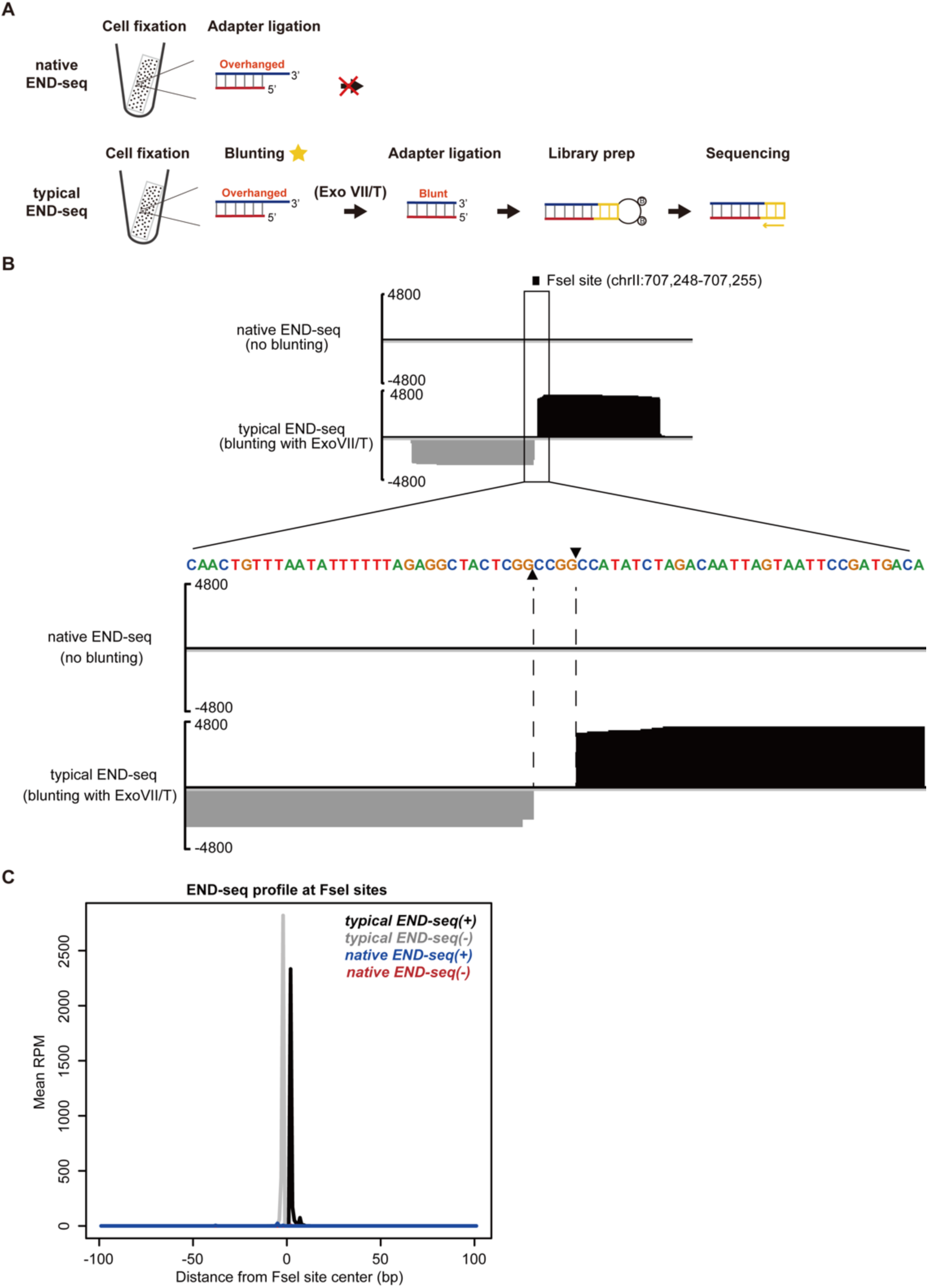
Sticky ends generated by FseI digestion cannot be captured by native END-seq, related to. Figure 3. (A) Schematic illustration showing the detection of sticky ends. Sticky ends cannot be ligated to hairpin adaptor for sequencing in native END-seq (top). Sticky ends blunted by Exo VII/T are ligated to hairpin adaptor for sequencing in typical END-seq (bottom). (B) Genomic DNA is digested *in vitro* by FseI, then blunted without (native END-seq) or with (typical END-seq) Exo VII/T. The restriction enzyme FseI generates 4-nt (5’-<u>CCGG</u>-3’) 3’-prochuding ends on genomic DNA. Top panel: DNA end generated by FseI digestion is enriched by typical END-seq, but not native END-seq. Bottom panel: zoomed in FseI site, reads at 3’ overhang (5‘-CCGG-3’) is absent in typical END-seq because of the blunting by Exo VII/T. (C) END-seq profile at the two sides of FseI sites. RPM, reads per million.

**Figure S6.**
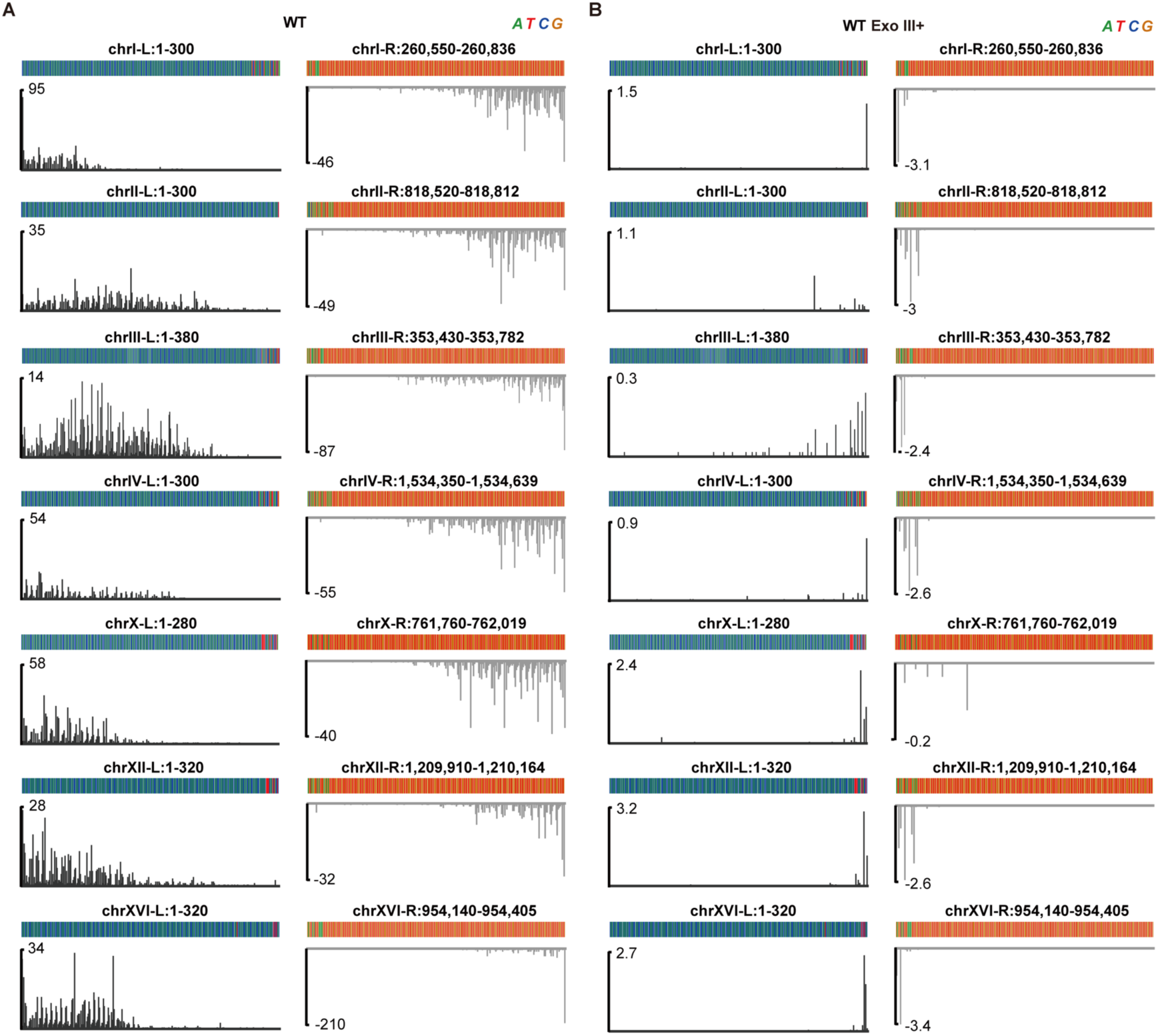
Native END-seq telomeric reads aligned to individual annotated telomeres, related to. Figure 3. (A and B) Representative alignments of telomeric reads to individual annotated telomere in the absence (A) and in the presence (B) of Exo III treatment (3’-5’ exonuclease for blunt end) in native END-seq.

**Figure S7.**
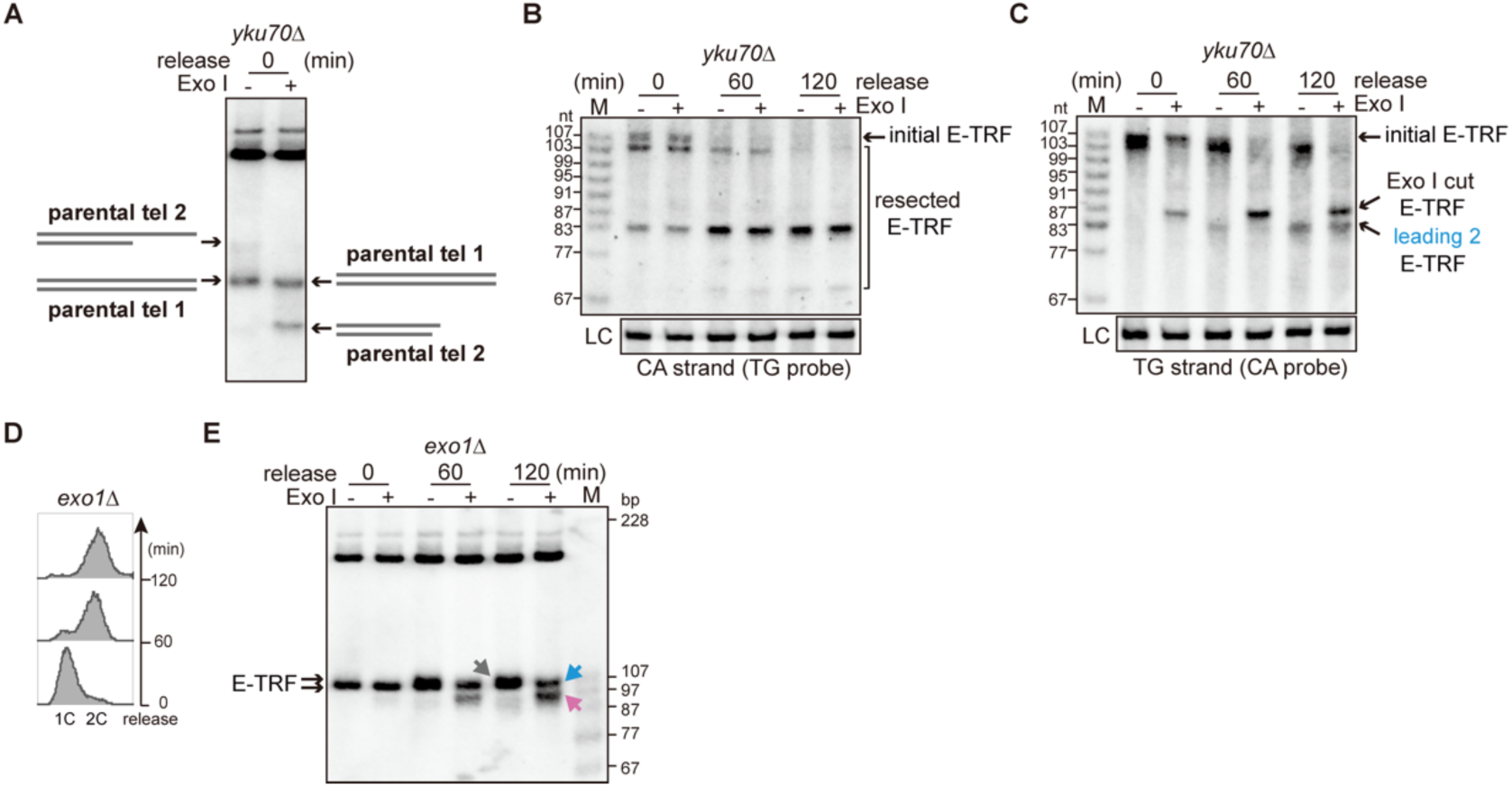
Both daughter telomeres are resected by nuclease in *yku70*Δ cells, related to. Figure 4. (A) Illustration of the E-TRF structures before replication in *yku70*Δ cells. The end structure of parental telomere is mainly divided into two types: parental tel 1 ends bluntly (refer to E-TRF 1 in Figure 3B, insensitive to Exo I treatment), while parental tel 2 ends with 24 nt overhang (refer to E-TRF 2 in Figure 3B, converts to 87 bp upon Exo I treatment), consistent with earlier results shown in Figure 1F. (B and C) Denatured Southern blotting analysis of the CA-strand (B) and TG-strand (C) during once replication in *yku70*Δ cells, hybridized with TG (B) and CA (C) probe, respectively. After replication, most of the CA strand was 83 nt and a few were ∼70 nt (B, 120 min). The TG strand was measured as 103 nt and 83 nt after replication, and the 103 nt product was converted to 87 nt when treated with Exo I (C, 120 min). (D) FACS analysis of DNA content in *exo1*Δ cells. (E) Southern blotting analysis of the 3’ overhang at both the leading and lagging strand telomere ends during once DNA replication in *exo1*Δ cells. Grey arrow: overlapped daughter telomeres, blue arrow: leading telomere, pink arrow: lagging telomere.

**Figure S8.**
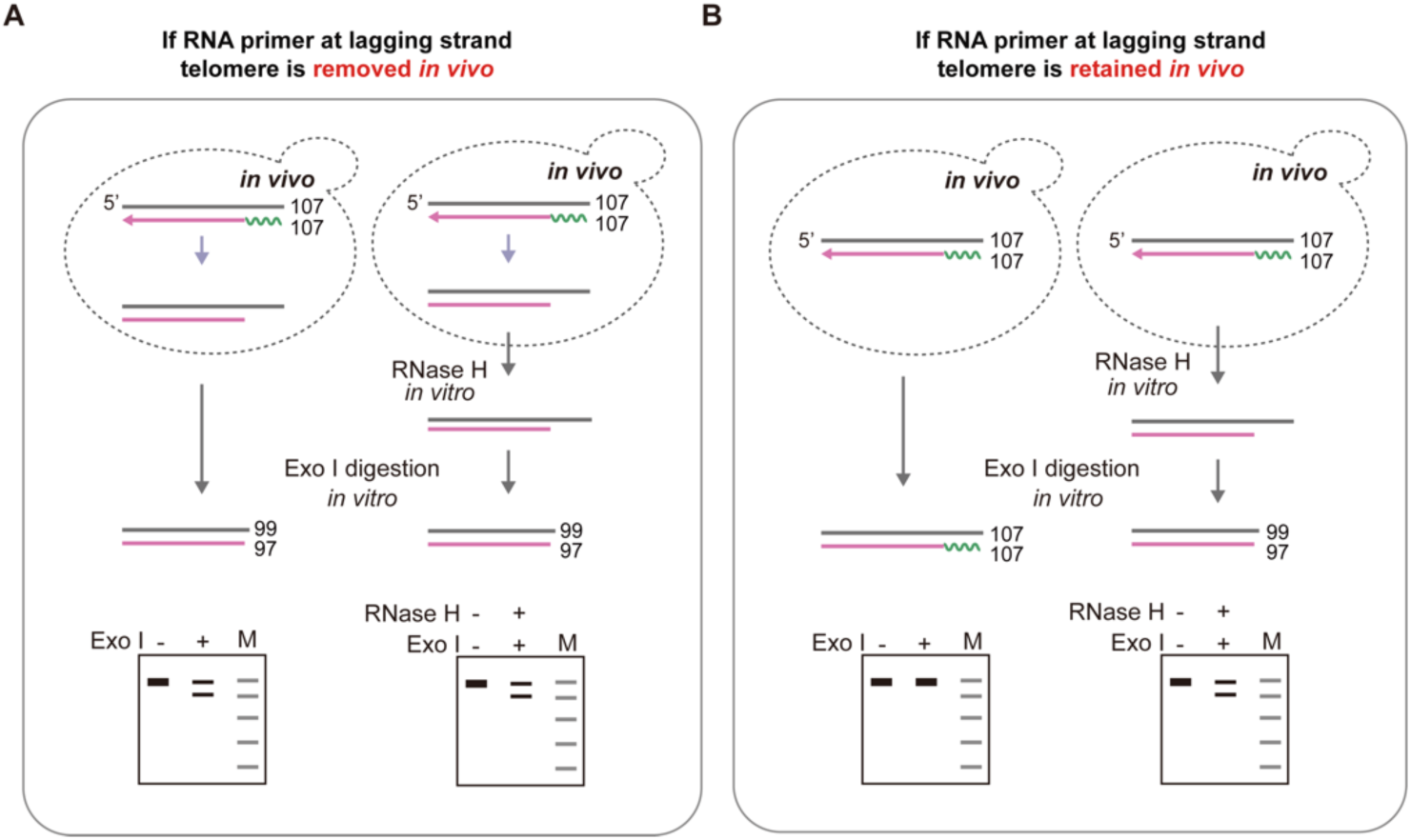
Schematic illustration of the RNA primer detection, related to. Figure 5. (A) *In vivo* RNA primer (green squiggly line) removal after replication leaves an overhang of ∼10 nt on lagging strand. The genomic DNA show similar sensitivity to *in vitro* Exo I digestion in the absence or presence of RNase H. (B) RNA primer (green squiggly line) is not removed after replication, the lagging strand ends with a blunt DNA-RNA primer hybrid. The genomic DNA is insensitive to *in vitro* Exo I digestion when not treated with RNase H (left panel), but becomes sensitive to *in vitro* Exo I digestion because of a 3’ overhang generated by RNase H (right panel).

**Figure S9.**
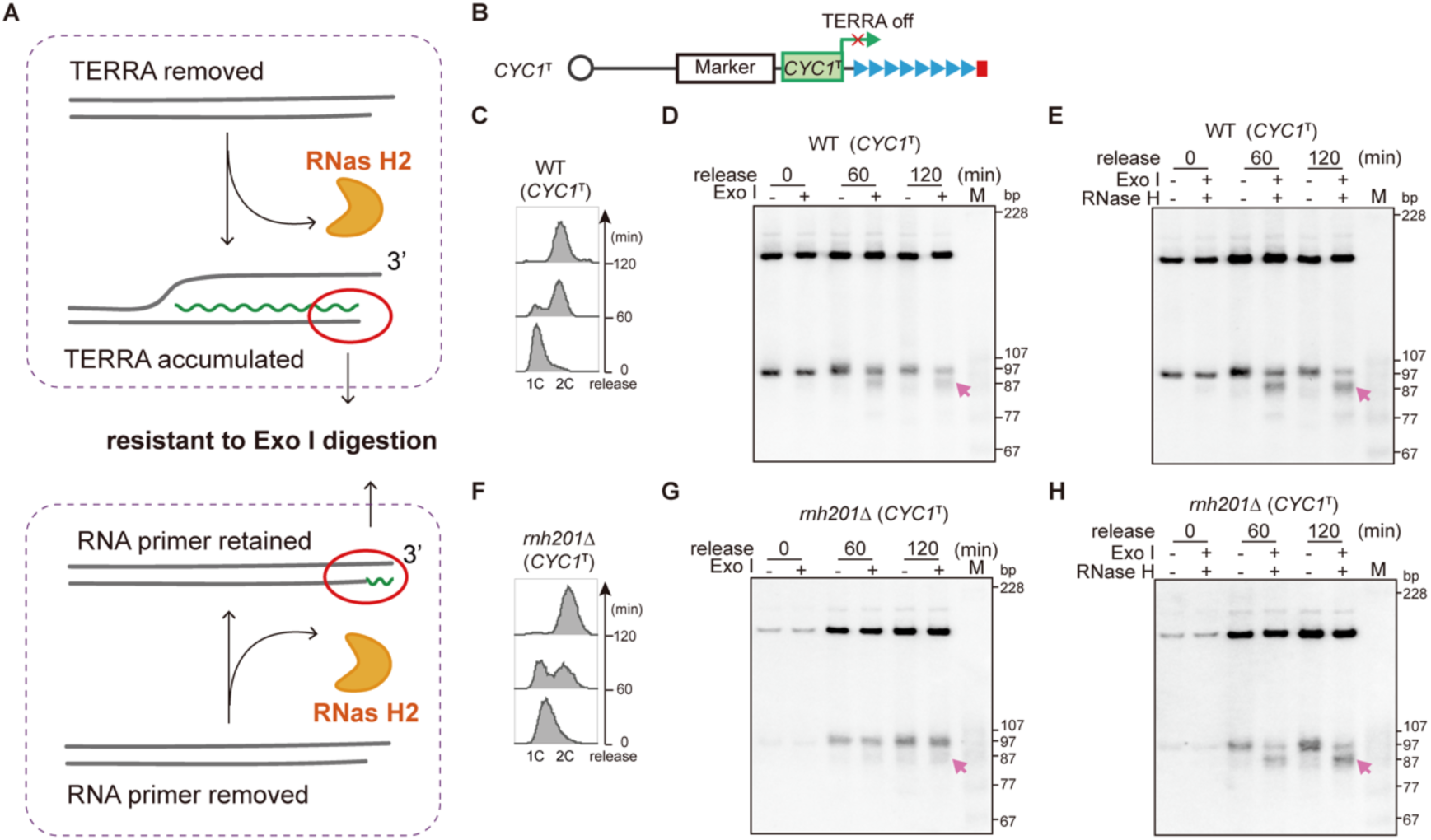
The DNA-RNA hybrid retained on lagging-strand telomere in *rnh201*Δ cells is DNA-RNA primer hybrid, related to. Figure 5. (A) Structure illustration of the accumulated TERRA DNA-RNA hybrid and DNA-RNA primer hybrid in absence of RNase H2. (B) Schematic illustration of the insertion of the *CYC1* terminator (*CYC1*^T^) at the upstream of the *de novo* telomere. (C and F) FACS analysis of DNA content in WT (*CYC1*^T^) (C) and *rnh201*Δ (*CYC1*^T^) (F) cells. (D, E, F and H) Southern blotting analysis of the 3’ overhang at both the leading and lagging strand telomere ends during once DNA replication in WT (*CYC1*^T^) (D, E) and *rnh201*Δ (*CYC1*^T^) (G, H) cells, pink arrow: Exo I trimmed lagging telomere. Genomic DNA is pretreated with RNase H as labeled.

**Figure S10.**
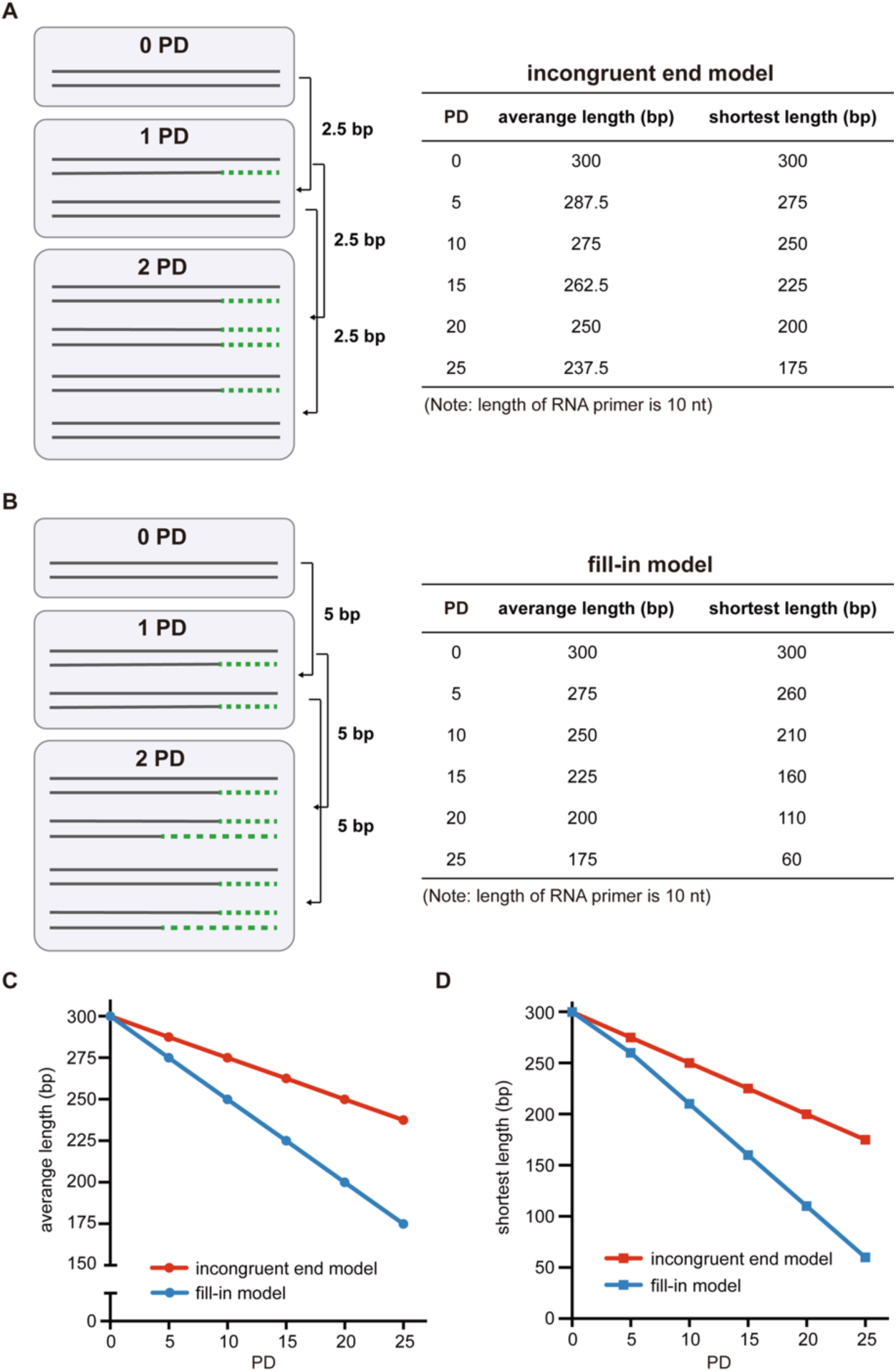
Schematic illustration of telomere shortening during consecutive cell divisions, related to. Figure 6. (A and B) Left panel, schematic illustration of telomere shortening for the first three rounds replication according to the incongruent end model (A) and fill-in model (B). Right panel, calculation of *de novo* telomere length during 25 population doublings according to the incongruent end model (A) and fill-in model (B). (C and D) Average telomere length (C) and shortest telomere length (D) during 25 population doublings (PD) according to the indicated models. Telomere shortening rate in fill-in model is faster than that in the incongruent end model.

**Figure S11.**
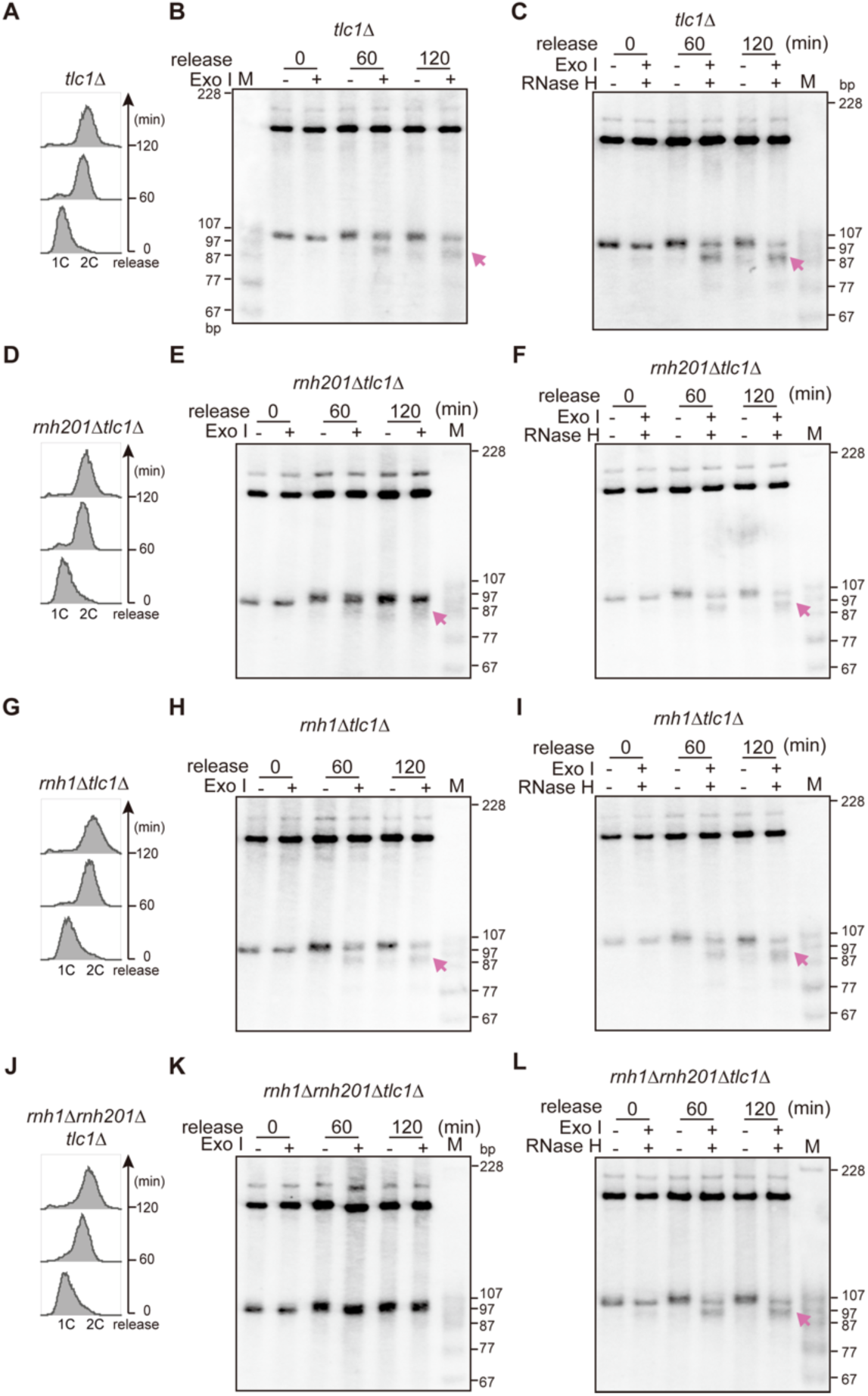
The RNA primer is retained on lagging telomere in absence of *RNH201* in telomerase null cells, related to. Figure 6. (A, D, G and J) FACS analysis of DNA content in *tlc1*Δ (A), *rnh201*Δ *tlc1*Δ (D), *rnh1*Δ *tlc1*Δ (G) and *rnh1*Δ *rnh201*Δ *tlc1*Δ (J) cells. (B, E, H and K) Southern blotting analysis of the 3’ overhang at both the leading and lagging strand telomere ends during once DNA replication in *tlc1*Δ (B), *rnh201*Δ *tlc1*Δ (E), *rnh1*Δ *tlc1*Δ (H) and *rnh1*Δ *rnh201*Δ *tlc1*Δ (K) cells, pink arrow: Exo I trimmed lagging telomere. (C, F, I and L) Southern blotting analysis of the 3’ overhang at both the leading and lagging strand telomere ends during once DNA replication in *tlc1*Δ (C), *rnh201*Δ *tlc1*Δ (F), *rnh1*Δ *tlc1*Δ (I) and *rnh1*Δ *rnh201*Δ *tlc1*Δ (L) cells, pink arrow: Exo I trimmed lagging telomere. Genomic DNA is pretreated with RNase H as labeled.

**Figure S12.**
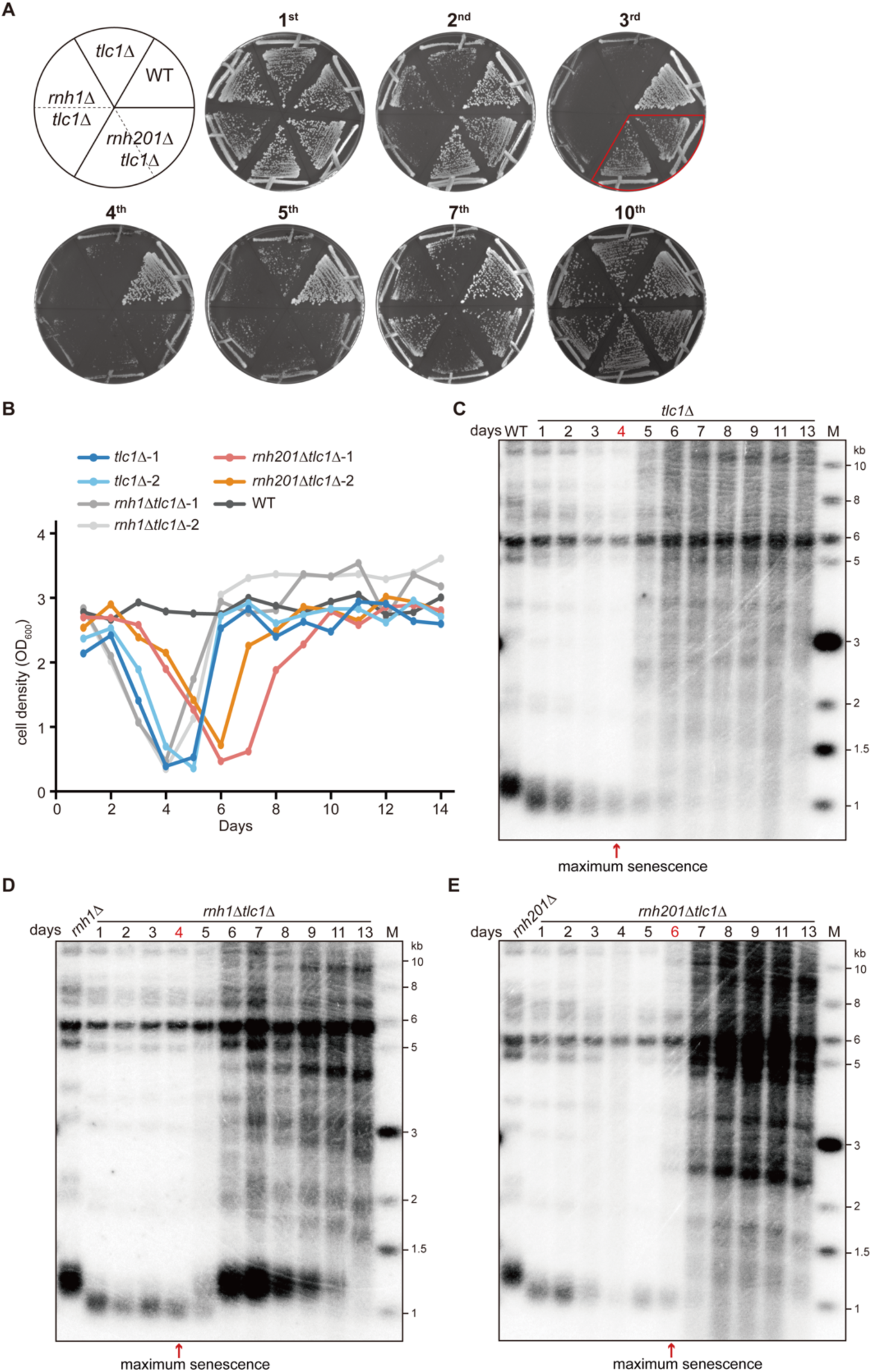
*RNH201* deletion delays cellular senescence and attenuates telomere shortening, related to. Figure 7. (A and B) The isogenic telomerase null mutants (*tlc1*Δ, *rnh1*Δ *tlc1*Δ and *rnh201*Δ *tlc1*Δ) were streaked on plate for successive 10 restreaks (A), or passaged in liquid media for 14 days (B) to monitor cellular senescence. (C-E) Southern blotting analysis of bulk telomeres in *tlc1*Δ (C), *rnh1*Δ *tlc1*Δ (D) and *rnh201*Δ *tlc1*Δ (E) cells that underwent senescence and survivor generation. Genomic DNA is digested with Xho I, and hybridized with a specific telomeric CA probe. The point of maximum senescence during passage is pointed by red arrow.

**Figure S13.**
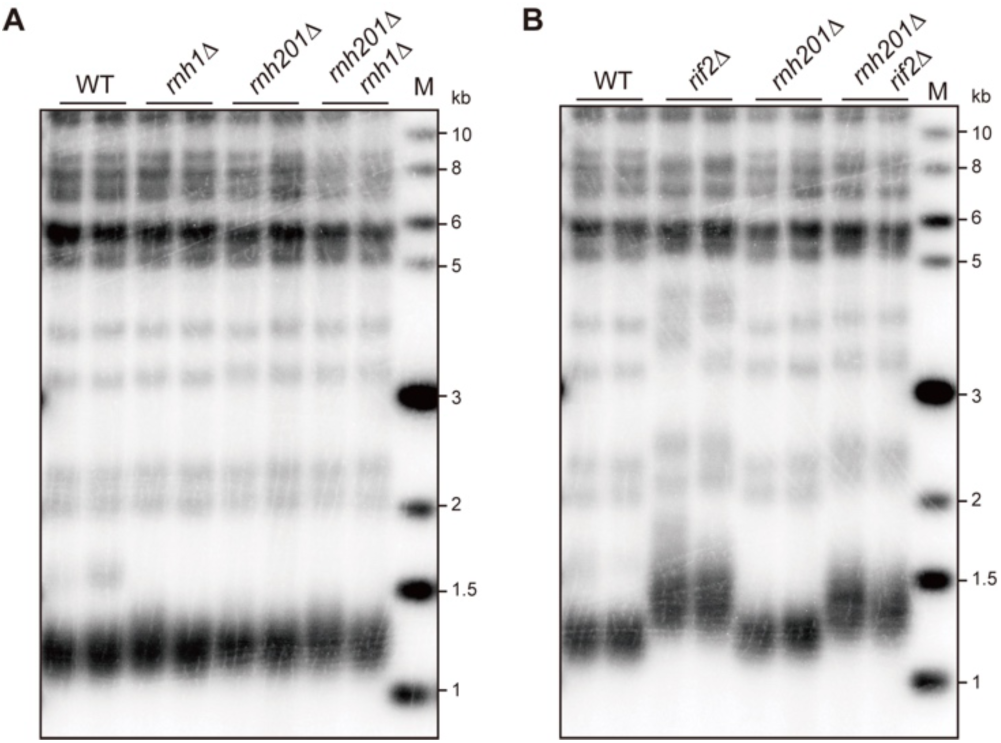
RNA primer retention has no effect on telomerase activity. (A) Southern blotting analysis of bulk telomere in RNase H mutants, genotypes of the strains are labeled on top, genomic DNA is digested with Xho I. (B) Southern blotting analysis of bulk telomere in *rif2*Δ cells with or without RNA primer retained on lagging telomere, genotypes of the strains are labeled on top, genomic DNA is digested with Xho I.

---

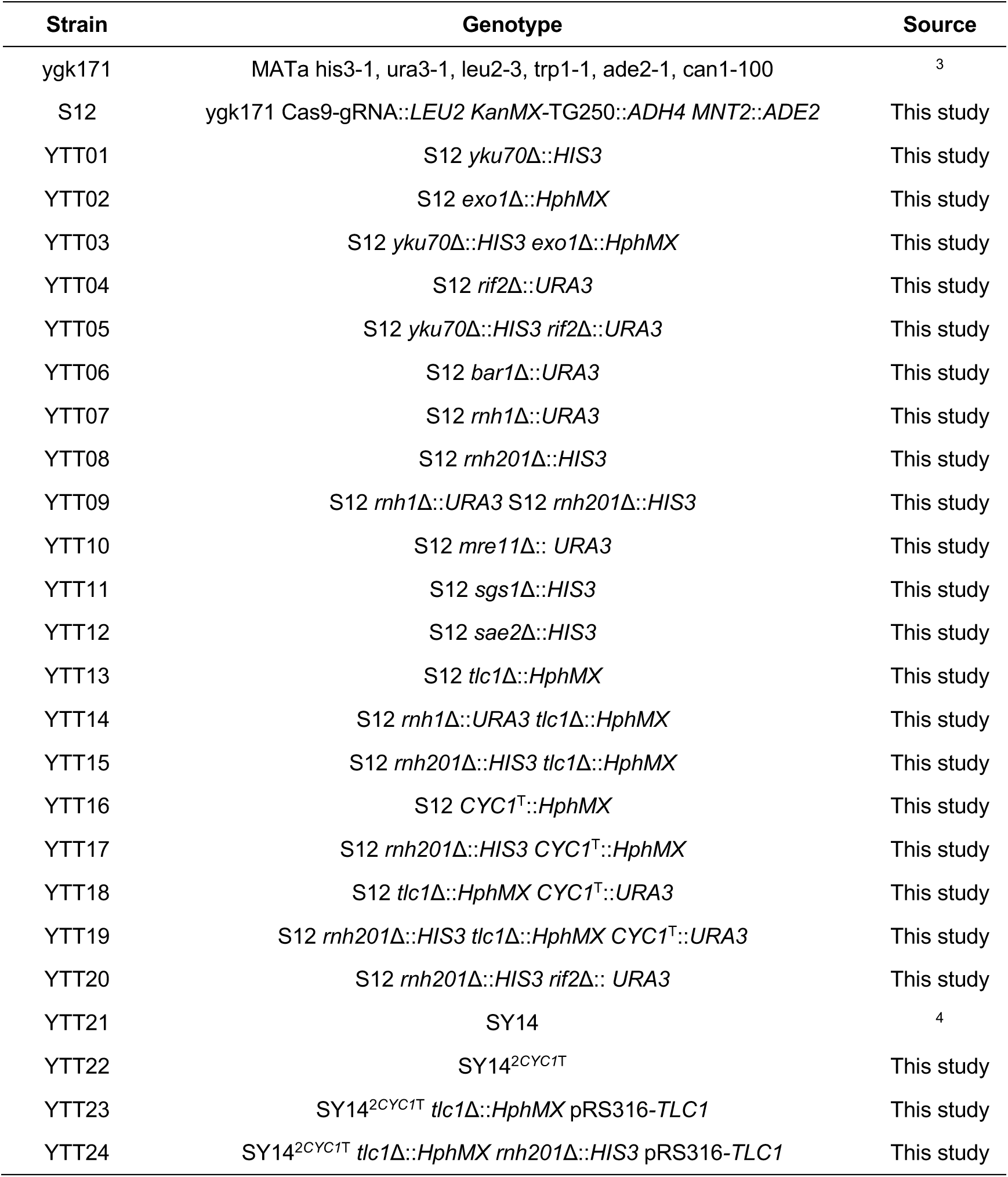
Supplementary information, Table S1. Strains used in this study.

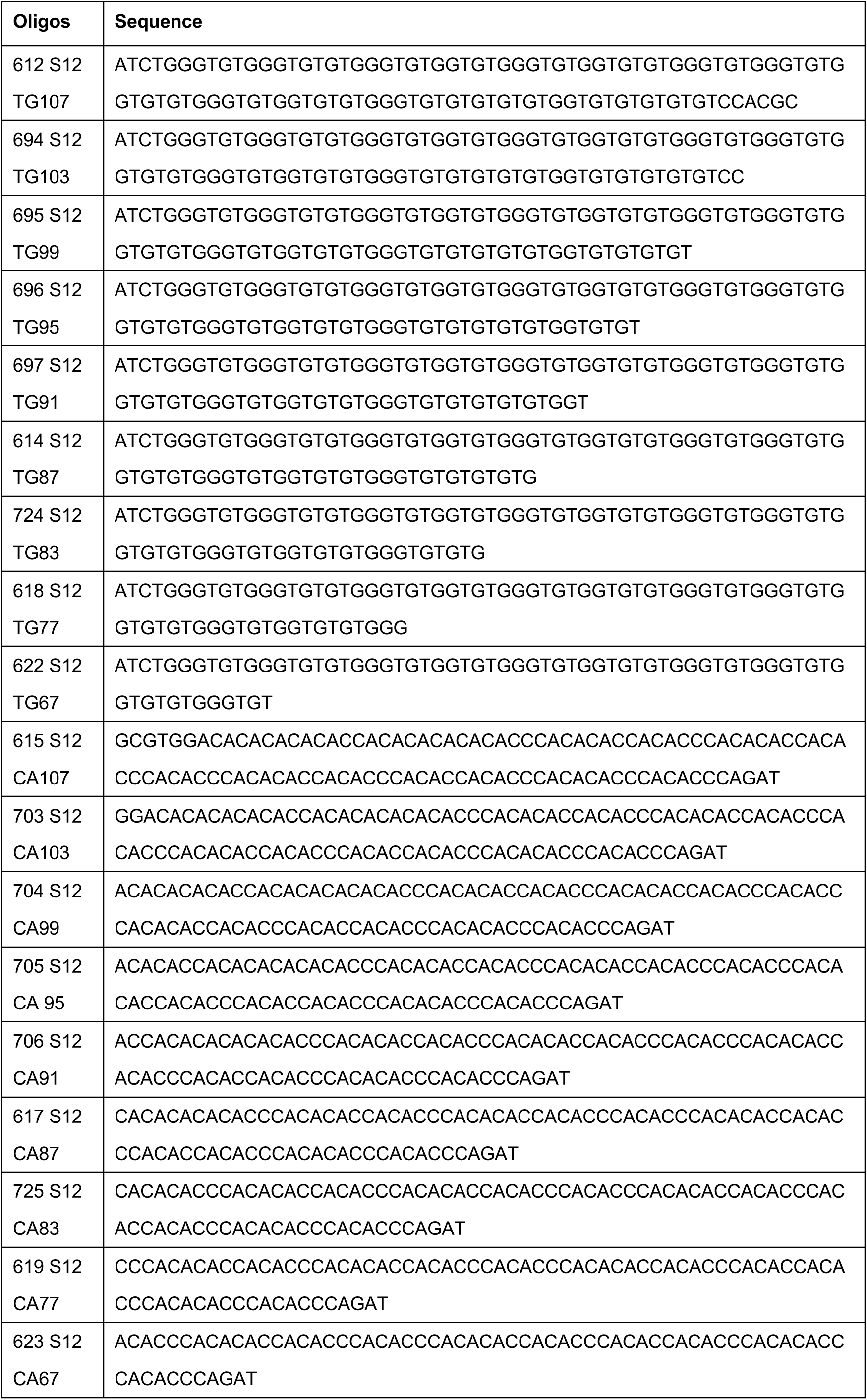
Supplementary information, Table S2. Oligonucleotide markers used in denatured PAGE.

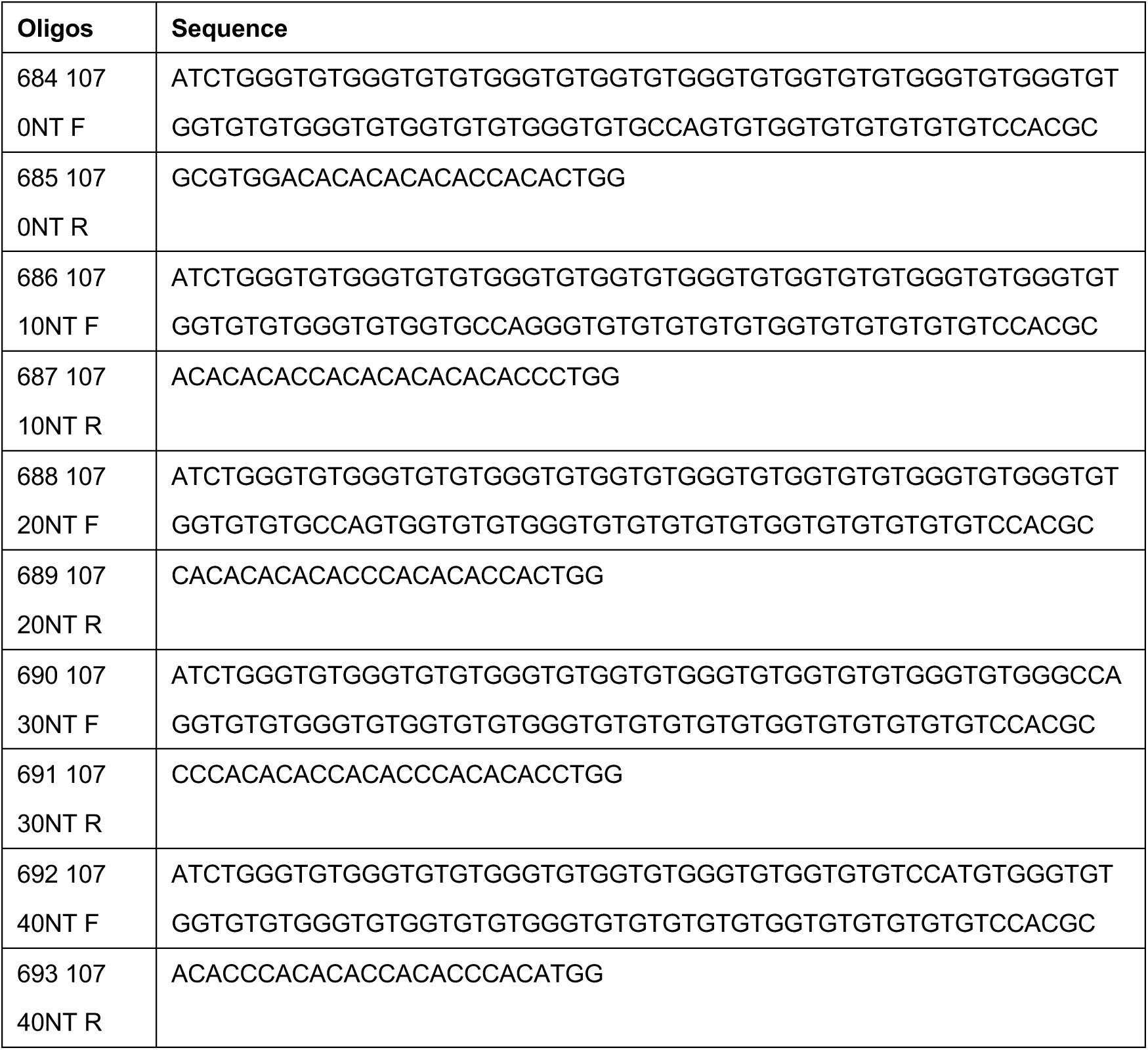
Supplementary information, Table S3. Oligonucleotide markers used in native PAGE.

Supplementary information, Table S4. Reagents used in this study.

**Supplementary information, Table S4. Reagents used in this study**
| REAGENT OR RESOURCE | SOURCE | IDENTIFIER |
| --- | --- | --- |
| **Propidium Iodide** | Sigma | P4170 |
| **Hydroxyurea** | Sigma | H8627 |
| **Nocodazole** | Sigma | M1404 |
| **Proteinase K** | Sigma | P6556 |
| **Nylon membrane** | Thermo Scientific | AM10104 |
| **RNase A** | Thermo Scientific | EN0531 |
| **EcoR V** | Thermo Scientific | FD0304 |
| **SuperSignal™ West Atto** | Thermo Scientific | A38556 |
| **Chemiluminescent Nucleic Acid**<br>**Detection Module Kit** | Thermo Scientific | 89880 |
| **BamH I** | Thermo Scientific | FD0054 |
| **Xho I** | Thermo Scientific | FD0694 |
| **dATP** | Thermo Scientific | 10216018 |
| **dNTP** | Thermo Scientific | 39025 |
| **Dynabeads MyOne C1** | Thermo Scientific | 65001 |
| **DynaMag™-2** | Thermo Scientific | 12321D |
| **Exo I** | NEB | M0293L |
| **RNase H** | NEB | M0297L |
| **Klenow Fragment (3'→5' exo-)** | NEB | M0212 |
| **Quick Ligase** | NEB | M2200L |
| **Buffer 2** | NEB | B7002S |
| **USER** | NEB | M5505S |
| **T4 Pollymerase** | NEB | M0203L |
| **β-Agarase I** | NEB | M0392S |
| **T4 Polynucleotide kinase** | NEB | M0201L |
| **Klenow fragment** | NEB | M0210L |
| **KAPA HiFi HotStart ReadyMix Kit** | ROCHE | 7958935001 |
| **VAHTS Multiplex Oligos Set 4 for**<br>**Illumina** | Vazyme | N321 |
| **CHEF Genomic DNA Plug Kits** | Bio-Rad | 170-3593 |
| **Electrophoresis mode** | Cytiva | SE260 |
| **Nuclease free water** | Beyotime | ST876 |
| **DNA low bind tube** | Eppendorf | 30108418 |
| **Wide bore tips** | AXYGEN | T250-WB CR |
| **Tips** | Brand | 732024 |
| **Probe** | Genscript | - |
| **Primer** | Azenta | - |
| **α-Factor** | Sangon | - |

## REFERENCES

1 Blackburn, E. H. Telomeres. Trends Biochem Sci 16, 378–381, doi:10.1016/0968-0004(91)90155-o (1991).

2 Pfeiffer, V. & Lingner, J. Replication of telomeres and the regulation of telomerase. Cold Spring Harb Perspect Biol 5, a010405, doi:10.1101/cshperspect.a010405 (2013).

3 Larrivée, M., LeBel, C. & Wellinger, R. J. The generation of proper constitutive G-tails on yeast telomeres is dependent on the MRX complex. Genes & development 18, 1391–1396 (2004).

4 Soudet, J., Jolivet, P. & Teixeira, M. T. Elucidation of the DNA end-replication problem in Saccharomyces cerevisiae. Mol Cell 53, 954–964, doi:10.1016/j.molcel.2014.02.030 (2014).

5 Gustafsson, C., Rhodin Edso, J. & Cohn, M. Rap1 binds single-stranded DNA at telomeric double-and single-stranded junctions and competes with Cdc13 protein. J Biol Chem 286, 45174–45185, doi:10.1074/jbc.M111.300517 (2011).

6 Wotton, D. & Shore, D. A novel Rap1p-interacting factor, Rif2p, cooperates with Rif1p to regulate telomere length in Saccharomyces cerevisiae. Genes Dev 11, 748–760, doi:10.1101/gad.11.6.748 (1997).

7 Ge, Y. et al. Structural insights into telomere protection and homeostasis regulation by yeast CST complex. Nat Struct Mol Biol 27, 752–762, doi:10.1038/s41594-020-0459-8 (2020).

8 Gravel, S., Larrivee, M., Labrecque, P. & Wellinger, R. J. Yeast Ku as a regulator of chromosomal DNA end structure. Science 280, 741–744, doi:10.1126/science.280.5364.741 (1998).

9 Wellinger, R. J. & Zakian, V. A. Everything you ever wanted to know about Saccharomyces cerevisiae telomeres: beginning to end. Genetics 191, 1073–1105, doi:10.1534/genetics.111.137851 (2012).

10 Wellinger, R. J. In the end, what’s the problem? Mol Cell 53, 855–856, doi:10.1016/j.molcel.2014.03.008 (2014).

11 Singer, M. S. & Gottschling, D. E. TLC1: template RNA component of Saccharomyces cerevisiae telomerase. *Science (New* York, N.Y.) 266, 404–409 (1994).

12 Lundblad, V. & Szostak, J. W. A mutant with a defect in telomere elongation leads to senescence in yeast. Cell 57, 633–643, doi:10.1016/0092-8674(89)90132-3 (1989).

13 Lundblad, V. & Blackburn, E. H. An alternative pathway for yeast telomere maintenance rescues est1-senescence. Cell 73, 347–360, doi:10.1016/0092-8674(93)90234-h (1993).

14 Zaug, A. J., Goodrich, K. J., Song, J. J., Sullivan, A. E. & Cech, T. R. Reconstitution of a telomeric replicon organized by CST. Nature 608, 819–825, doi:10.1038/s41586-022-04930-8 (2022).

15 Chen, H. et al. Structural Insights into Yeast Telomerase Recruitment to Telomeres. Cell 172, 331–343 e313, doi:10.1016/j.cell.2017.12.008 (2018).

16 Hoerr, R. E., Ngo, K. & Friedman, K. L. When the Ends Justify the Means: Regulation of Telomere Addition at Double-Strand Breaks in Yeast. Front Cell Dev Biol 9, 655377, doi:10.3389/fcell.2021.655377 (2021).

17 Fu, X. H. et al. Telomere recombination preferentially occurs at short telomeres in telomerase-null type II survivors. PLoS One 9, e90644, doi:10.1371/journal.pone.0090644 (2014).

18 Marcand, S., Brevet, V. & Gilson, E. Progressive cis-inhibition of telomerase upon telomere elongation. EMBO J 18, 3509–3519, doi:10.1093/emboj/18.12.3509 (1999).

19 Kuchta, R. D., Reid, B. & Chang, L. M. DNA primase. Processivity and the primase to polymerase alpha activity switch. Journal of Biological Chemistry 265, 16158–16165, doi:10.1016/s0021-9258(17)46202-8 (1990).

20 Sun, H. et al. Okazaki fragment maturation: DNA flap dynamics for cell proliferation and survival. Trends Cell Biol 33, 221–234, doi:10.1016/j.tcb.2022.06.014 (2023).

21 Nguyen, T. A. et al. Analysis of subunit assembly and function of the Saccharomyces cerevisiae RNase H2 complex. FEBS J 278, 4927–4942, doi:10.1111/j.1742-4658.2011.08394.x (2011).

22 Lockhart, A. et al. RNase H1 and H2 Are Differentially Regulated to Process RNA-DNA Hybrids. Cell Rep 29, 2890–2900 e2895, doi:10.1016/j.celrep.2019.10.108 (2019).

23 Luke, B. et al. The Rat1p 5’ to 3’ exonuclease degrades telomeric repeat-containing RNA and promotes telomere elongation in Saccharomyces cerevisiae. Mol Cell 32, 465–477, doi:10.1016/j.molcel.2008.10.019 (2008).

24 Balk, B. et al. Telomeric RNA-DNA hybrids affect telomere-length dynamics and senescence. Nat Struct Mol Biol 20, 1199–1205, doi:10.1038/nsmb.2662 (2013).

25 Misino, S., Busch, A., Wagner, C. B., Bento, F. & Luke, B. TERRA increases at short telomeres in yeast survivors and regulates survivor associated senescence (SAS). Nucleic Acids Res 50, 12829–12843, doi:10.1093/nar/gkac1125 (2022).

26 Huang, L. et al. Role of calf RTH-1 nuclease in removal of 5’-ribonucleotides during Okazaki fragment processing. Biochemistry 35, 9266–9277, doi:10.1021/bi9603074 (1996).

27 Murante, R. S., Henricksen, L. A. & Bambara, R. A. Junction ribonuclease: an activity in Okazaki fragment processing. Proc Natl Acad Sci U S A 95, 2244–2249, doi:10.1073/pnas.95.5.2244 (1998).

28 Lai, T. P., Wright, W. E. & Shay, J. W. Comparison of telomere length measurement methods. Philos Trans R Soc Lond B Biol Sci 373, doi:10.1098/rstb.2016.0451 (2018).

29 Shampay, J. & Blackburn, E. H. Generation of telomere-length heterogeneity in Saccharomyces cerevisiae. Proc Natl Acad Sci U S A 85, 534–538, doi:10.1073/pnas.85.2.534 (1988).

30 Ribeyre, C. & Shore, D. Regulation of telomere addition at DNA double-strand breaks. Chromosoma 122, 159–173, doi:10.1007/s00412-013-0404-2 (2013).

31 Diede, S. J. & Gottschling, D. E. Telomerase-mediated telomere addition in vivo requires DNA primase and DNA polymerases alpha and delta. Cell 99, 723–733 (1999).

32 Negrini, S., Ribaud, V., Bianchi, A. & Shore, D. DNA breaks are masked by multiple Rap1 binding in yeast: implications for telomere capping and telomerase regulation. Genes Dev 21, 292–302, doi:10.1101/gad.400907 (2007).

33 Bonetti, D., Martina, M., Clerici, M., Lucchini, G. & Longhese, M. P. Multiple pathways regulate 3’ overhang generation at S. cerevisiae telomeres. Mol Cell 35, 70–81, doi:10.1016/j.molcel.2009.05.015 (2009).

34 Eldridge, A. M., Halsey, W. A. & Wuttke, D. S. Identification of the determinants for the specific recognition of single-strand telomeric DNA by Cdc13. Biochemistry 45, 871–879, doi:10.1021/bi0512703 (2006).

35 Vignais, M. L., Huet, J., Buhler, J. M. & Sentenac, A. Contacts between the factor TUF and RPG sequences. Journal of Biological Chemistry 265, 14669–14674, doi:10.1016/s0021-9258(18)77354-7 (1990).

36 Bertuch, A. A. & Lundblad, V. The Ku heterodimer performs separable activities at double-strand breaks and chromosome termini. Mol Cell Biol 23, 8202–8215, doi:10.1128/MCB.23.22.8202-8215.2003 (2003).

37 Vodenicharov, M. D., Laterreur, N. & Wellinger, R. J. Telomere capping in non-dividing yeast cells requires Yku and Rap1. EMBO J 29, 3007–3019, doi:10.1038/emboj.2010.155 (2010).

38 Bonetti, D. et al. Shelterin-like proteins and Yku inhibit nucleolytic processing of Saccharomyces cerevisiae telomeres. PLoS Genet 6, e1000966, doi:10.1371/journal.pgen.1000966 (2010).

39 Hirano, Y., Fukunaga, K. & Sugimoto, K. Rif1 and rif2 inhibit localization of tel1 to DNA ends. Mol Cell 33, 312–322, doi:10.1016/j.molcel.2008.12.027 (2009).

40 Ribeyre, C. & Shore, D. Anticheckpoint pathways at telomeres in yeast. Nat Struct Mol Biol 19, 307–313, doi:10.1038/nsmb.2225 (2012).

41 Maringele, L. & Lydall, D. EXO1-dependent single-stranded DNA at telomeres activates subsets of DNA damage and spindle checkpoint pathways in budding yeast yku70Delta mutants. Genes Dev 16, 1919–1933, doi:10.1101/gad.225102 (2002).

42 Cannavo, E. & Cejka, P. Sae2 promotes dsDNA endonuclease activity within Mre11-Rad50-Xrs2 to resect DNA breaks. Nature 514, 122–125, doi:10.1038/nature13771 (2014).

43 Hardy, J., Churikov, D., Geli, V. & Simon, M. N. Sgs1 and Sae2 promote telomere replication by limiting accumulation of ssDNA. Nat Commun 5, 5004, doi:10.1038/ncomms6004 (2014).

44 Chow, T. T., Zhao, Y., Mak, S. S., Shay, J. W. & Wright, W. E. Early and late steps in telomere overhang processing in normal human cells: the position of the final RNA primer drives telomere shortening. Genes Dev 26, 1167–1178, doi:10.1101/gad.187211.112 (2012).

45 Wu, P., Takai, H. & de Lange, T. Telomeric 3’ overhangs derive from resection by Exo1 and Apollo and fill-in by POT1b-associated CST. Cell 150, 39–52, doi:10.1016/j.cell.2012.05.026 (2012).

46 Canela, A. et al. DNA Breaks and End Resection Measured Genome-wide by End Sequencing. Mol Cell 63, 898–911, doi:10.1016/j.molcel.2016.06.034 (2016).

47 Wong, N., John, S., Nussenzweig, A. & Canela, A. END-seq: An Unbiased, High-Resolution, and Genome-Wide Approach to Map DNA Double-Strand Breaks and Resection in Human Cells. Methods Mol Biol 2153, 9–31, doi:10.1007/978-1-0716-0644-5_2 (2021).

48 Wu, W. et al. Neuronal enhancers are hotspots for DNA single-strand break repair. Nature 593, 440–444, doi:10.1038/s41586-021-03468-5 (2021).

49 Hughes, T. R., Weilbaecher, R. G., Walterscheid, M. & Lundblad, V. Identification of the single-strand telomeric DNA binding domain of the Saccharomyces cerevisiae Cdc13 protein. Proc Natl Acad Sci U S A 97, 6457–6462, doi:10.1073/pnas.97.12.6457 (2000).

50 Porro, A., Feuerhahn, S., Reichenbach, P. & Lingner, J. Molecular dissection of telomeric repeat-containing RNA biogenesis unveils the presence of distinct and multiple regulatory pathways. Mol Cell Biol 30, 4808–4817, doi:10.1128/MCB.00460-10 (2010).

51 Graf, M. et al. Telomere Length Determines TERRA and R-Loop Regulation through the Cell Cycle. Cell 170, 72–85 e14, doi:10.1016/j.cell.2017.06.006 (2017).

52 Shao, Y. et al. Creating a functional single-chromosome yeast. Nature 560, 331–335, doi:10.1038/s41586-018-0382-x (2018).

53 Cristofari, G. & Lingner, J. Telomere length homeostasis requires that telomerase levels are limiting. EMBO J 25, 565–574, doi:10.1038/sj.emboj.7600952 (2006).

54 Chang, M., Arneric, M. & Lingner, J. Telomerase repeat addition processivity is increased at critically short telomeres in a Tel1-dependent manner in Saccharomyces cerevisiae. Genes Dev 21, 2485–2494, doi:10.1101/gad.1588807 (2007).

55 Lundblad, V. Telomere end processing: unexpected complexity at the end game. Genes Dev 26, 1123–1127, doi:10.1101/gad.195339.112 (2012).

56 Sfeir, A. J., Chai, W., Shay, J. W. & Wright, W. E. Telomere-end processing the terminal nucleotides of human chromosomes. Mol Cell 18, 131–138, doi:10.1016/j.molcel.2005.02.035 (2005).

57 Cech, T. R. Beginning to understand the end of the chromosome. Cell 116, 273–279, doi:10.1016/s0092-8674(04)00038-8 (2004).

58 Marcomini, I. et al. Asymmetric Processing of DNA Ends at a Double-Strand Break Leads to Unconstrained Dynamics and Ectopic Translocation. Cell Rep 24, 2614–2628 e2614, doi:10.1016/j.celrep.2018.07.102 (2018).

59 Cai, S. W. & de Lange, T. CST-Polalpha/Primase: the second telomere maintenance machine. Genes Dev 37, 555–569, doi:10.1101/gad.350479.123 (2023).

60 Emerson, C. H. et al. Ku DNA End-Binding Activity Promotes Repair Fidelity and Influences End-Processing During Nonhomologous End-Joining in Saccharomyces cerevisiae. Genetics 209, 115–128, doi:10.1534/genetics.117.300672 (2018).

61 Wright, W. E., Tesmer, V. M., Huffman, K. E., Levene, S. D. & Shay, J. W. Normal human chromosomes have long G-rich telomeric overhangs at one end. Genes Dev 11, 2801–2809, doi:10.1101/gad.11.21.2801 (1997).

62 Riha, K., McKnight, T. D., Fajkus, J., Vyskot, B. & Shippen, D. E. Analysis of the G-overhang structures on plant telomeres: evidence for two distinct telomere architectures. Plant J 23, 633–641, doi:10.1046/j.1365-313x.2000.00831.x (2000).

63 Ferreira, M. G., Miller, K. M. & Cooper, J. P. Indecent exposure: when telomeres become uncapped. Mol Cell 13, 7–18, doi:10.1016/s1097-2765(03)00531-8 (2004).

64 Greider, C. W. Regulating telomere length from the inside out: the replication fork model. Genes Dev 30, 1483–1491, doi:10.1101/gad.280578.116 (2016).

65 Sfeir, A. & de Lange, T. Removal of shelterin reveals the telomere end-protection problem. Science 336, 593–597, doi:10.1126/science.1218498 (2012).

66 de Lange, T. Shelterin-Mediated Telomere Protection. Annu Rev Genet 52, 223–247, doi:10.1146/annurev-genet-032918-021921 (2018).

67 Burgers, P. M. J. & Kunkel, T. A. Eukaryotic DNA Replication Fork. Annu Rev Biochem 86, 417–438, doi:10.1146/annurev-biochem-061516-044709 (2017).

68 Qiu, J., Qian, Y., Frank, P., Wintersberger, U. & Shen, B. Saccharomyces cerevisiae RNase H(35) functions in RNA primer removal during lagging-strand DNA synthesis, most efficiently in cooperation with Rad27 nuclease. Mol Cell Biol 19, 8361–8371, doi:10.1128/MCB.19.12.8361 (1999).

69 Wu, T. J. et al. Sequential loading of Saccharomyces cerevisiae Ku and Cdc13p to telomeres. J Biol Chem 284, 12801–12808, doi:10.1074/jbc.M809131200 (2009).

70 Krasner, D. S., Daley, J. M., Sung, P. & Niu, H. Interplay between Ku and Replication Protein A in the Restriction of Exo1-mediated DNA Break End Resection. J Biol Chem 290, 18806–18816, doi:10.1074/jbc.M115.660191 (2015).

71 Takai, H., Aria, V., Borges, P., Yeeles, J. T. P. & de Lange, T. CST-polymerase alpha-primase solves a second telomere end-replication problem. Nature, doi:10.1038/s41586-024-07137-1 (2024).

72 Ohki, R., Tsurimoto, T. & Ishikawa, F. In vitro reconstitution of the end replication problem. Mol Cell Biol 21, 5753–5766, doi:10.1128/MCB.21.17.5753-5766.2001 (2001).

73 Ray, A. & Runge, K. W. The yeast telomere length counting machinery is sensitive to sequences at the telomere-nontelomere junction. Mol Cell Biol 19, 31–45, doi:10.1128/MCB.19.1.31 (1999).

74 McGee, J. S. et al. Reduced Rif2 and lack of Mec1 target short telomeres for elongation rather than double-strand break repair. Nat Struct Mol Biol 17, 1438–1445, doi:10.1038/nsmb.1947 (2010).

75 Bonnell, E., Pasquier, E. & Wellinger, R. J. Telomere Replication: Solving Multiple End Replication Problems. Front Cell Dev Biol 9, 668171, doi:10.3389/fcell.2021.668171 (2021).

76 Olovnikov, A. M. [Principle of marginotomy in template synthesis of polynucleotides]. Dokl Akad Nauk SSSR 201, 1496–1499 (1971).

77 Audoynaud, C. et al. RNA:DNA hybrids from Okazaki fragments contribute to establish the Ku-mediated barrier to replication-fork degradation. Mol Cell 83, 1061–1074 e1066, doi:10.1016/j.molcel.2023.02.008 (2023).

78 Liu, G. et al. RPA-mediated recruitment of Bre1 couples histone H2B ubiquitination to DNA replication and repair. Proc Natl Acad Sci U S A 118, doi:10.1073/pnas.2017497118 (2021).

79 Rohner, S., Gasser, S. M. & Meister, P. Modules for cloning-free chromatin tagging in Saccharomyces cerevisae. Yeast 25, 235–239, doi:10.1002/yea.1580 (2008).

80 Sikorski, R. S. & Hieter, P. A system of shuttle vectors and yeast host strains designed for efficient manipulation of DNA in Saccharomyces cerevisiae. Genetics 122, 19–27, doi:10.1093/genetics/122.1.19 (1989).

81 Rosebrock, A. P. Synchronization and Arrest of the Budding Yeast Cell Cycle Using Chemical and Genetic Methods. Cold Spring Harb Protoc 2017, doi:10.1101/pdb.prot088724 (2017).

82 Wu, Z. J. et al. Cdc13 is predominant over Stn1 and Ten1 in preventing chromosome end fusions. Elife 9, doi:10.7554/eLife.53144 (2020).

83 Haase, S. B. & Lew, D. J. Flow cytometric analysis of DNA content in budding yeast. Methods Enzymol 283, 322–332, doi:10.1016/s0076-6879(97)83026-1 (1997).

84 Liu, L. et al. Tracking break-induced replication shows that it stalls at roadblocks. Nature 590, 655–659, doi:10.1038/s41586-020-03172-w (2021).

## REFERENCES

1 Soudet, J., Jolivet, P. & Teixeira, M. T. Elucidation of the DNA end-replication problem in Saccharomyces cerevisiae. Mol Cell 53, 954–964, doi:10.1016/j.molcel.2014.02.030 (2014).

2 Wellinger, R. J. In the end, what’s the problem? Mol Cell 53, 855–856, doi:10.1016/j.molcel.2014.03.008 (2014).

3 Liu, G. et al. RPA-mediated recruitment of Bre1 couples histone H2B ubiquitination to DNA replication and repair. Proc Natl Acad Sci U S A 118, doi:10.1073/pnas.2017497118 (2021).

4 Shao, Y. et al. Creating a functional single-chromosome yeast. Nature 560, 331–335, doi:10.1038/s41586-018-0382-x (2018).

